# Malnutrition drives infection susceptibility and dysregulated myelopoiesis that persists after refeeding intervention

**DOI:** 10.1101/2024.08.19.608575

**Authors:** Alisa Sukhina, Clemence Queriault, Saptarshi Roy, Elise Hall, Kelly Rome, Muskaan Aggarwal, Elizabeth Nunn, Ashley Weiss, Janet Nguyen, F Chris Bennett, Will Bailis

**Affiliations:** Department of Pathology and Laboratory Medicine, Children’s Hospital of Philadelphia; Philadelphia, PA, USA; Department of Pathology and Laboratory Medicine, Perelman School of Medicine, University of Pennsylvania; Philadelphia, PA, USA; Department of Physiology, Perelman School of Medicine at the University of Pennsylvania, Philadelphia, PA, USA; Department of Psychiatry, Perelman School of Medicine, University of Pennsylvania, Philadelphia, PA, USA; Division of Neurology, Children’s Hospital of Philadelphia, Philadelphia, PA, USA

## Abstract

Undernutrition is one of the largest persistent global health crises, with nearly 1 billion people facing severe food insecurity. Infectious disease represents the main underlying cause of morbidity and mortality for malnourished individuals, with infection during malnutrition representing the leading cause of childhood mortality worldwide. In the face of this complex challenge, simple refeeding protocols have remained the primary treatment strategy. Although an association between undernutrition and infection susceptibility has been appreciated for over a century, the underlying mechanisms remain poorly understood and the extent to which refeeding intervention is sufficient to reverse nutritionally acquired immunodeficiency is unclear. Here we investigate how malnutrition leads to immune dysfunction and the ability of refeeding to repair it. We find that chronic malnutrition induced through prolonged dietary restriction (40% reduction in food intake) severely impairs the ability of mice to control a sub-lethal *Listeria monocytogenes* infection. Malnourished mice exhibit blunted immune cell expansion, impaired immune function, and accelerated contraction prior to pathogen clearance. While this defect is global, we find that myelopoiesis is uniquely impacted, resulting in reduced neutrophil and monocyte numbers prior to and post-infection. Upon refeeding, we observe that mice recover body mass, size, cellularity across all major immune organs, the capacity to undergo normal immune cell expansion in response to infection, and a restoration in T cell responses. Despite this broad improvement, refed mice remain susceptible to *Listeria* infection, uncoupling global lymphoid atrophy from immunodeficiency. We find peripheral neutrophil and monocyte numbers fail to fully recover and refed mice are unable to undergo normal emergency myelopoiesis. Altogether, this work identifies dysregulated myelopoiesis as a link between prior nutritional state and immunocompetency. We believe these findings raise the possibility that exposure to food scarcity should be treated as an immunologic variable, even post-recovery, with considerations for how patient medical history and public health policy.

## Introduction

Almost 500 million adults and over 200 million children are affected by undernutrition worldwide, with over half of all childhood deaths linked to undernutrition (United Nations, 2021; UNICEF, 1998). In the past 5 years, these numbers worsened due to economic instability ushered by COVID-19 pandemic, and they are projected to further increase throughout the impending climate crisis (Mbow et al., 2019; United Nations, 2022). In patients with undernutrition, the largest contributor to morbidity and mortality is infectious disease (Fan et al., 2022; Rice et al., 2000). It has been demonstrated that undernutrition alone puts patients, especially children, at a higher risk of developing long-lasting disability or dying from an infection (Antwi 2011; Bhargava, 2016; Bourke et al., 2016; Caulfield et al., 2004; Rice et al., 2000; Schlaudecker et al., 2011; Sinha et al., 2021). In addition to increased susceptibility to infection, chronically undernourished individuals have been reported to have impaired barrier function, atrophy of immune organs, and less effective responses to vaccines (Bhattacharjee et al., 2021; Bourke et al., 2016; Collins and Belkaid, 2020; Prendergast, 2015). In keeping with this, undernutrition is the leading cause of secondary immunodeficiency in the world (Chinen and Shearer, 2010). Considering the geographical overlap between areas with high prevalence of undernutrition and infectious disease and the persistent nature of the global undernutrition crisis, it is critical to investigate how undernutrition causes immunodeficiency and contributes to poor infection outcomes (Mbow et al., 2019; Murray et al., 1996; Pelletier, 1994; Roberts, 2017; Rohr et al., 2019; Sinha et al., 2021; UNICEF-WHO-World Bank JME Working Group, 2021).

Despite the first link between undernutrition and poor infection outcomes being posited over a century ago, the cellular mechanisms underlying undernutrition-induced immunodeficiency remain poorly resolved (Bourke et al., 2019, 2016; Hess, 1932; Newsholme, 1908). The primary explanation given for this dysfunction has centered on defects in lymphocyte biology. Patient data and animal studies suggest that T, B, and NK cells are reduced in the periphery of patients with undernutrition and experimental animals (Campbell et al., 2020; Cason et al., 1986; Contreras et al., 2018; Howard et al., 1999; Nájera et al., 2004; Saha et al., 1977; Schattner et al., 1990; Yang et al., 2009). Moreover, animal models of short-term fasting and protein deficiency have been found to impair T cell expansion, cytokine production, and memory recall responses (Chatraw et al., 2008; Iyer et al., 2012; Mangheri et al., 1992; Procaccini et al., 2012; Saucillo et al, 2014; Taylor et al., 2013). While the contribution of lymphocyte dysfunction to nutritionally acquired immunodeficiency is well established, the impact of prolonged undernutrition on other immune cell populations and the role they play in disease susceptibility is not well understood.

Prior studies have reported disparate impacts of undernutrition on immunity and infection, with some finding dietary or caloric restriction enhances inflammation or immunity, while others observing a loss in immune function (Bhattacharjee et al., 2021; Campbell et al., 2020; Chatraw et al., 2008; Collins et al., 2019; Han et al., 2023; Hasegawa et al., 2012; Iyer et al., 2012; Palma et al., 2021; Pena-cruz et al, 1989; Piccio et al., 2008; Starr et al., 2016; Sun et al., 2001; Taylor et al., 2013). These discrepancies likely result from whether animals are subjected to caloric restriction alone or if there is also corresponding limitation in key micro- and macronutrients and whether the restriction in food intake is administered as a short-term fast or is sustained (Cerqueira and Kowaltowski, 2010; Contreras et al., 2018; Meydani et al., 2016; Palma et al., 2021; Yan et al., 2021). To this end, caloric and nutrient restriction diets significantly differ in weight loss patterns, physiology, and nutritional state (Cerqueira and Kowaltowski, 2010). While both diets are associated with reduced inflammation, caloric restriction is understood to support both lifespan and health span, in stark contrast to chronic undernutrition (Collins and Belkaid, 2020; Contreras et al., 2018; Goldberg et al., 2015; Green et al., 2021; Hasegawa et al., 2012; Howard et al., 1999; Jordan et al., 2019; Piccio et al., 2008; Spadaro et al., 2022; Sun et al., 2001; Yang et al., 2009).

Beyond our mechanistic understanding of nutritionally acquired immunodeficiency, there is a limited knowledge on whether this dysfunction is reversible (Contreras et al., 2018; Schattner et al., 1990; Vaisman et al., 2004). Because current treatment guidelines for patients with undernutrition utilize refeeding protocols that focus on weight gain as the indicator of recovery, the effects of refeeding on the immune system are understudied (Ashworth et al., 2003). Moreover, whether prior exposure to malnutrition has durable effects on the ability to control infection following refeeding remains unknown.

Here, we investigate the impact of nutritionally acquired immunodeficiency on the immune responses to *Listeria monocytogenes* infection and the ability of refeeding intervention to restore immune competency. We employ a chronic murine dietary restriction model and demonstrate that it effectively recapitulates the hallmarks of human undernutrition. We find that chronic undernutrition results in a specific atrophy of immune organs that is not mirrored in essential organs, such as the liver and kidney. This loss of lymphoid tissue is accompanied by a broad reduction in both innate and adaptive immune cell compartments. Undernourished mice fail to control sub-lethal *L. monocytogenes* infection, either succumbing to disease or failing to clear pathogen long-term. We find that while these mice undergo the initial phase of immune cell expansion following infection, they fail to sustain it and the response rapidly undergoes contraction. Accordingly, we observe that T cells in these mice display muted expansion, accelerated contraction, and impaired effector function. We further find that undernutrition selectively impairs steady-state and emergency myelopoiesis, resulting in reduced neutrophil and monocyte abundance prior to and post-infection. Finally, we demonstrate that while refeeding protocols are sufficient to restore body mass, growth, and reverse global lymphoid atrophy, refed mice remain more susceptible to infection even months after recovering size and peripheral immune cell numbers. In contrast to the ability of refeeding to reverse the effects of undernutrition on lymphoid numbers, we go on to show refed mice display impaired emergency myelopoiesis as well as neutrophil and monocyte abundance. Altogether, our work identifies dysregulated myelopoiesis as a major factor contributing to increased susceptibility to infection during chronic undernutrition. Furthermore, we for the first time demonstrate that refeeding protocols are not sufficient to reverse defects in the ability of mice to control infection or dysregulated myelopoiesis, despite outwardly displaying a recovery from malnutrition. We believe these findings have important implications for global public health policy and medical standards of care not only for treating people actively experiencing chronic malnutrition, but also for individuals who have experienced and recovered from food scarcity as part of their life history.

## Results

### Sustained dietary restriction recapitulates the hallmarks of nutritionally acquired immunodeficiency

To investigate the relationship of undernutrition and immunodeficiency, we began by adopting a faithful model of chronic malnutrition. In patients, undernutrition is associated with: (1) a consistent weight loss of 10% or more compared to the age-appropriate body weight that occurs in under 6 months and persists for months to years; (2) stunting of growth; and (3) a change in body composition associated with weight loss (Institute of Medicine, 2000; Maleta, 2006; Martins et al., 2011). While short-term fasting and macronutrient or caloric restriction models have been previously used, we elected to employ a 40% reduced diet (40RD) model, which provides both the weight loss as well as micro- and macronutrient deficiency over extended periods of time (Cerqueira and Kowaltowski, 2010). For the 40RD system, the average food intake of C57Bl6 mice was measured at baseline consumption. After a week of baseline measurements, mice were randomly assigned to either the *ad libitum* fed (AL) or the 40RD group. 40% of the average food intake by weight was removed from the diet of 40RD mice (Fig. 1a). Animal weight, body length, and body condition score (BCS) were then regularly surveyed to measure wasting, stunting, and body composition, respectively. Highlighting the ability of the model to reliably generate a state of moderate undernutrition, 40RD mice consistently lost 10% of their initial body weight (IBW) in 4 weeks (Fig. 1b) and achieved an average 20% loss of IBW relative to the AL group (Fig. 1c). At the same time, BCSs decreased significantly in 40RD mice compared to the AL group over the course of 4 weeks (Fig. 1d). This decrease in weight and conditioning was further accompanied by delayed growth and stunting in 40RD mice, as measured by body length (Fig. 1e).

**Figure 1.**
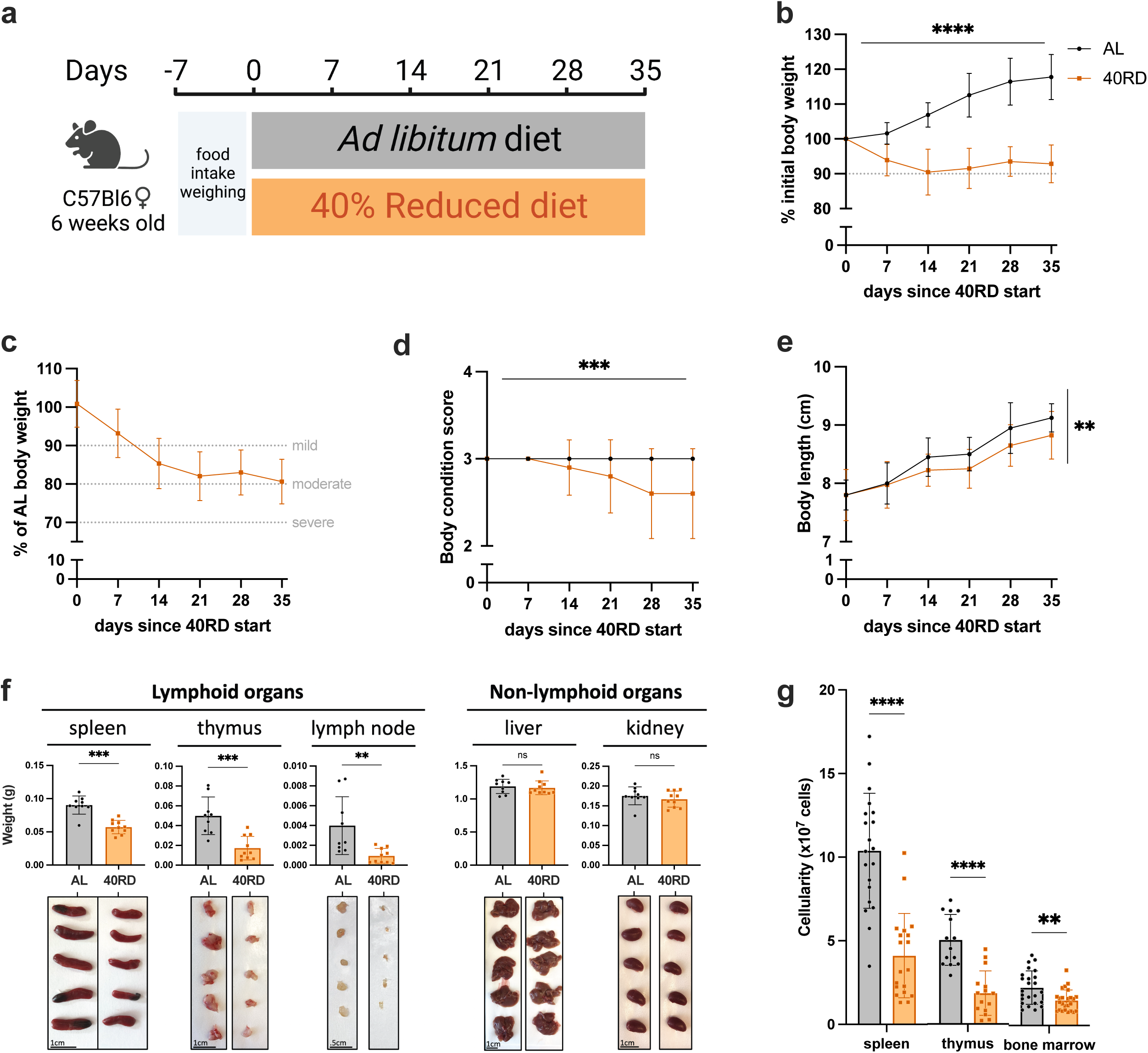
Sustained dietary restriction recapitulates the hallmarks of nutritionally acquired immunodeficiency. (a) Schematic of experimental design for 40% reduced diet (40RD, orange) in comparison to control *ad libitum* (AL, black) diet. (b) Body weight of AL and 40RD mice as a percentage of initial body weight over time (n=50). The dotted line represents 10% of initial body weight lost. (c) Body weight of 40RD mice as a percentage of age matched average AL body weight over time (n=50). Each dotted line represents clinical designations of undernutrition severity. (d) Body condition score of AL and 40RD mice over time (n=10). (e) Body length of AL and 40RD mice over time, measured from the nose tip to the base of the tail (n=10). (f) Comparative weights of AL and 40RD lymphoid and non-lymphoid tissues (n=10) with representative photos of the corresponding organs. Scale bars 1 cm (0.5 cm for lymph nodes). (g) Total live cell counts for whole spleen (n=15), thymus (n=15), and bone marrow (n=10). Statistics: (b-g) Plotted as mean ± SD; (b,d) simple linear regression with slope comparisons; (e) simple linear regression with elevation comparison; and (f,g) two-tailed Mann-Whitney test.

Beyond these gross changes in body condition, patients with undernutrition also exhibit severe lymphoid organ atrophy (Beisel, 1996; Bourke et al., 2016b). In keeping with this, we observed a reduction in spleen, thymus, and lymph node size and mass in mice given the 40RD diet, while organs such as the liver and kidney were unchanged, suggesting a specific effect of malnutrition on the immune system (Fig. 1f). We observed a corresponding reduction in the cellularity of the spleen and thymus, with a more modest loss of cellularity observed in the bone marrow (Fig. 1g). Collectively, our findings demonstrate that the 40RD model reproduces the lymphoid atrophy found in chronically patients with undernutrition as well as other models of undernutrition in rodents (Beisel, 1996; Bourke et al., 2016b; Cason et al., 1986; Contreras et al., 2018; Yang et al., 2009).

### Chronic malnutrition results in a failure to control sub-lethal L. monocytogenes infection

We next sought to investigate how malnutrition impacted immune responses to infection. While the link between poor infection outcomes and undernutrition has been extensively documented, the immunologic mechanisms resulting in poor infection resolution remain unknown (Bourke et al., 2019, 2016b; Dubos, 1955; Ishikawa et al., 2012; Rice et al., 2000b). To address this, we turned to the Ovalbumin expressing *Listeria monocytogenes* (Lm-Ova) system, a classic model of bacterial infection that permits the tracking of adaptive immunity, innate immunity, and bacterial burden. Both 40RD and AL mice were infected with a sublethal dose of Lm-Ova and then infection progression, pathogen clearance, and immune responses were assessed at 5, 8, and 14 days post-infection (Fig. 2a). Whereas all mice in the AL group survived, nearly half of the 40RD mice became moribund and required euthanasia (Fig. 2b). Even amongst surviving mice, the 40RD group displayed higher clinical scores than the AL group throughout the course of infection (Fig. 2c). Consistent with this, 40RD mice failed to clear bacteria at the rate of the AL group, with some mice exhibiting persistent infection (Fig. 2d). Amongst the AL group, 48.6% and 86.7% of the mice were able to clear bacteria by day 5 and day 8 post-infection, respectively, with all mice resolving infection by day 14. In contrast, 40RD mice exhibited delayed clearance kinetics with only a third of the mice becoming pathogen-free by day 14 (Fig. 2d).

**Figure 2.**
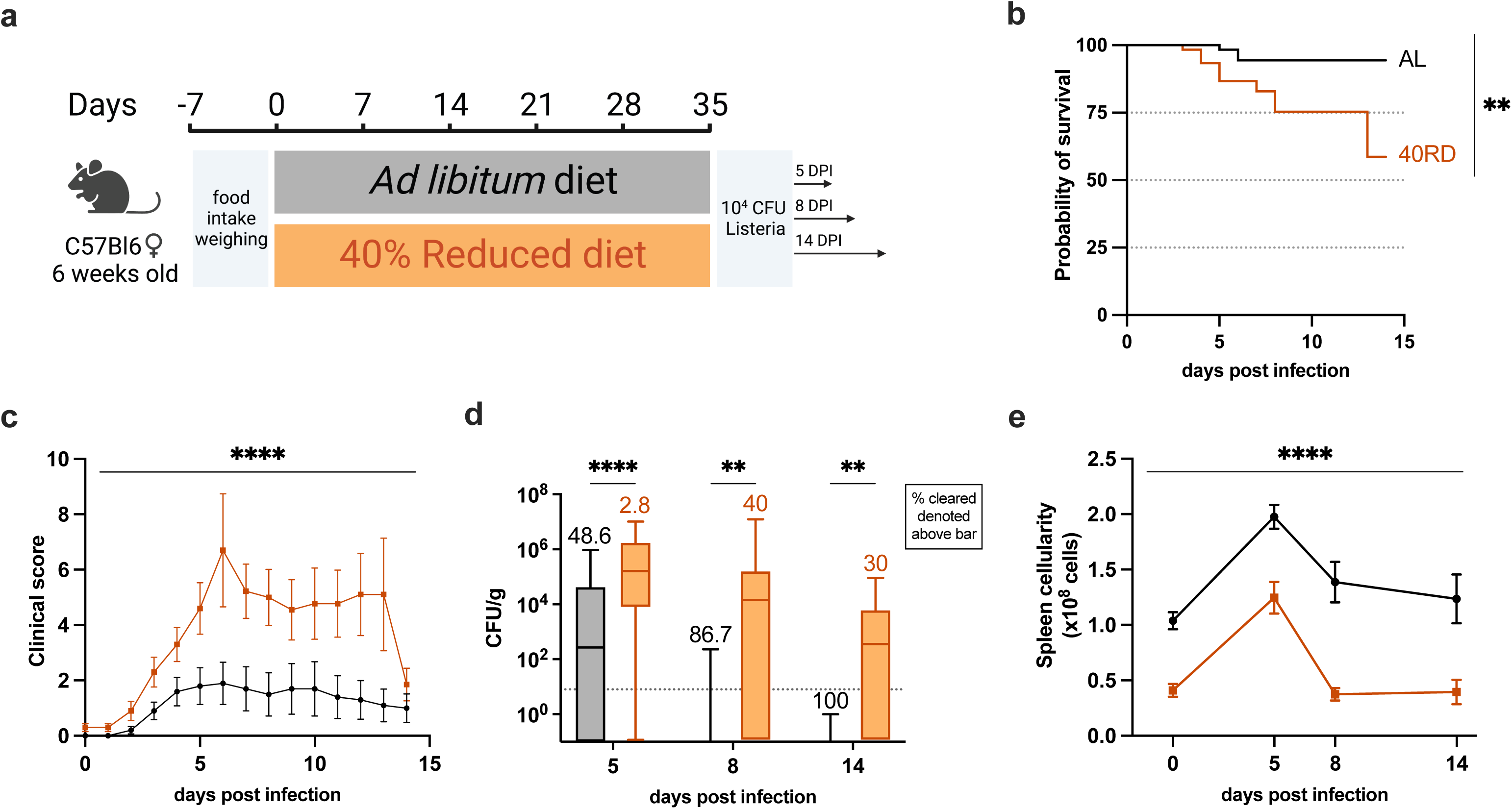
Chronic malnutrition results in a failure to control sub-lethal *L. monocytogenes* infection. (a) Schematic of Lm infection (10^4^ CFUs per mouse) experimental design in AL (orange) and 40RD (black) mice. Mice were maintained on the corresponding diet throughout the course of the infection. (b) Probability of survival for infected AL and 40RD mice over time. The curves represent pooled data from 3 experimental groups: 5DPI (n=25), 8DPI (n=15), and 14DPI (n=10). Statistics done via log-rank test. (c) Clinical score for infected AL and 40RD mice over time from 14DPI group. Plotted as mean ± SEM; statistics done via mixed-effect two-way ANOVA analysis. (d) Pathogen burden in liver tissue of AL and 40RD mice. Percentage of mice that cleared the pathogen on a given day is represented as numbers above corresponding bars. The dotted line represents the limit of detection. Plotted as box and min to max whiskers; statistics done via two-tailed Mann-Whitney test for each time point. (e) Total splenocyte counts for infected AL and 40RD mice over time. Uninfected spleen cell counts same as used for Figure 1g. Plotted as mean ± SEM; statistics done via mixed-effect two-way ANOVA analysis.

These observations prompted us to ask whether 40RD mice could mount a normal immune response to infection. To begin addressing this, we examined immune cell expansion in AL and 40RD mice throughout the course of infection. We found that while splenocyte numbers increased at early time points following infection in 40RD mice, the magnitude and duration of this expansion was significantly lower compared to AL controls (Fig. 2e). Thus, chronic malnutrition is sufficient to not only induce immune atrophy, but further abrogates the capacity of mice to mount and sustain an immune response to infection.

### Chronic malnutrition diminishes T cell expansion and function while accelerating contraction during infection

Considering the broad defects observed in infected, chronically undernourished mice, we next aimed to investigate the effects malnutrition had on specific immune cell populations during this response. As malnourished patients are known to exhibit the reduction in lymphocyte numbers, we assessed whether 40RD mice exhibited signs of lymphopenia following infection (Cason et al., 1986; Nájera et al., 2004; Saha et al., 1977). We observed that both prior to and after infection, 40RD mice displayed reduced numbers of B cells, CD4 T cells, and CD8 T cells compared to controls (Supplemental 1). Despite this overall loss in lymphocyte number, the relative frequency of each population was either unchanged or elevated, indicating that while malnutrition leads to a global reduction in immune cell numbers, lymphocytes are less impacted than other immune cell populations (Supplemental 1).

Considering the role CD8 T cells play in resolving intracellular bacterial infections and the failure of 40RD mice to clear infection at later time points, we next examined whether the kinetics of T cell expansion and contraction were similarly impacted. To do so, we tracked T cell function throughout the course of infection, on day 5 (early response), day 8 (peak response), and day 14 (contraction phase) post-infection (Qiu et al., 2018; Zenewicz and Shen, 2007). We found that the antigen-experienced (CD44^+^) and short-lived effector (CD127^low^KLRG1^high^) CD8^+^ T cell numbers were significantly reduced throughout the infection (Fig. 3a,b). These effector-like populations expanded minimally during early and peak response and contracted quickly by day 14, prior to pathogen clearance in 40RD mice. In addition to these defects in expansion/contraction dynamics, we further tested whether chronic malnutrition also impaired T cells function. We thus evaluated both the frequency and per-cell cytokine producing capacity of T cells at day 8 post-infection. We observed that T cells from undernourished mice displayed both a decreased frequency of cytokine producing cells and a decreased capacity for producing cytokines on a per-cell basis (Fig. 3c-e). Altogether, these findings suggest that impaired T cell function along with insufficient T cell expansion and premature contraction prior to pathogen clearance contribute to the failure of chronically undernourished mice in controlling *L. monocytogenes* infection.

**Figure 3.**
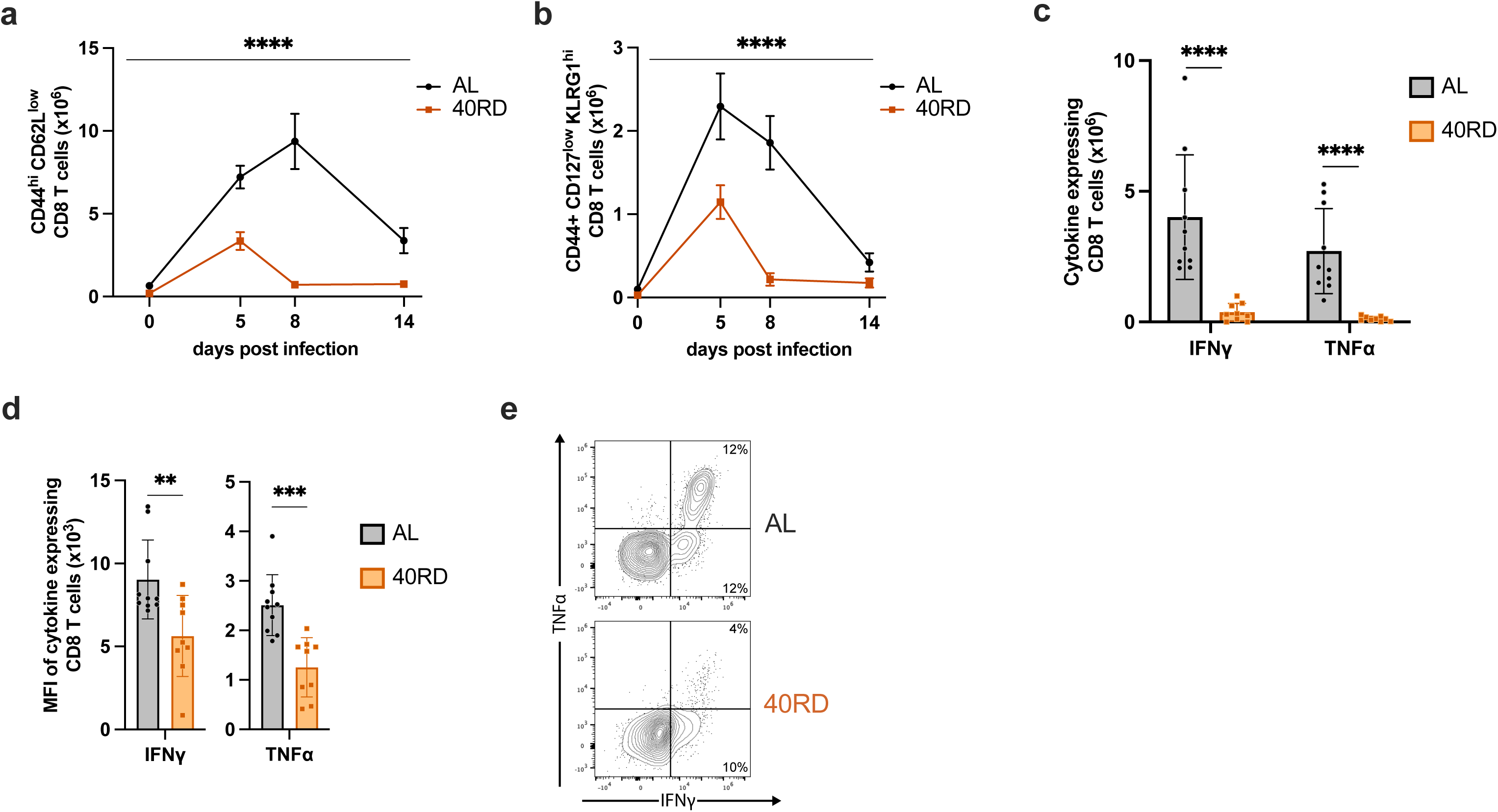
Chronic malnutrition diminishes T cell expansion and function while accelerating contraction during infection. AL and 40RD mice were infected with Lm at 10^4^ CFUs per mouse. Splenocytes were counted and flow cytometry was performed at days 0, 5, 8, and 14 post-infection to evaluate the total cell number of (a) antigen-experienced CD8^+^ T cells gated on live single CD8^+^ CD44^hi^ CD62L^low^ and (b) short lived effector cells further gated on KLRG1^hi^ CD127^low^. Plotted as mean ± SEM; statistics done via mixed-effect two-way ANOVA analysis. Splenocytes from AL (n=10) and 40RD (n=10) mice were harvest at day 8 post-infection and stimulated ex vivo with OVA peptide for 6 hours. Intracellular flow cytometry was performed to quantify (c) total cell number of and (d) mean fluorescence intensity (MFI) of antigen-experienced CD8^+^ T cells expressing IFNγ and TNFα. Plotted as mean ± SD; statistics done via two-tailed Mann-Whitney test for each cytokine. (e) Representative flow cytometry data of (c,d), with average frequencies shown.

### Chronically malnourished mice display dysregulated myelopoiesis

The observation that 40RD mice exhibited elevated pathogen burden at early time points prior to the peak of the adaptive immune response between days 5 to 8 prompted us to investigate whether innate immune responses were similarly impaired. While total bone marrow cellularity was equivalent between 40RD and AL groups, examination of specific innate immune populations in bone marrow revealed that steady-state neutrophil and monocyte levels were significantly lower in undernourished mice, both in absolute numbers and relative abundance (Fig. 1g & Fig. 4a-d; Supplemental 2). In keeping with this, we observed that splenic neutrophil and monocyte abundance were significantly lower in 40RD mice at steady-state and underwent impaired expansion following infection, with splenic monocyte frequency elevated at steady state and comparable to control mice post-infection (Fig. 4b-d; Supplemental 2). These steady-state changes could be observed early after mice were placed on the 40RD diet, with total peripheral blood cellularity and neutrophil abundance significantly reduced by 1 week post-dietary restriction (Supplemental 2e,f).

**Figure 4.**
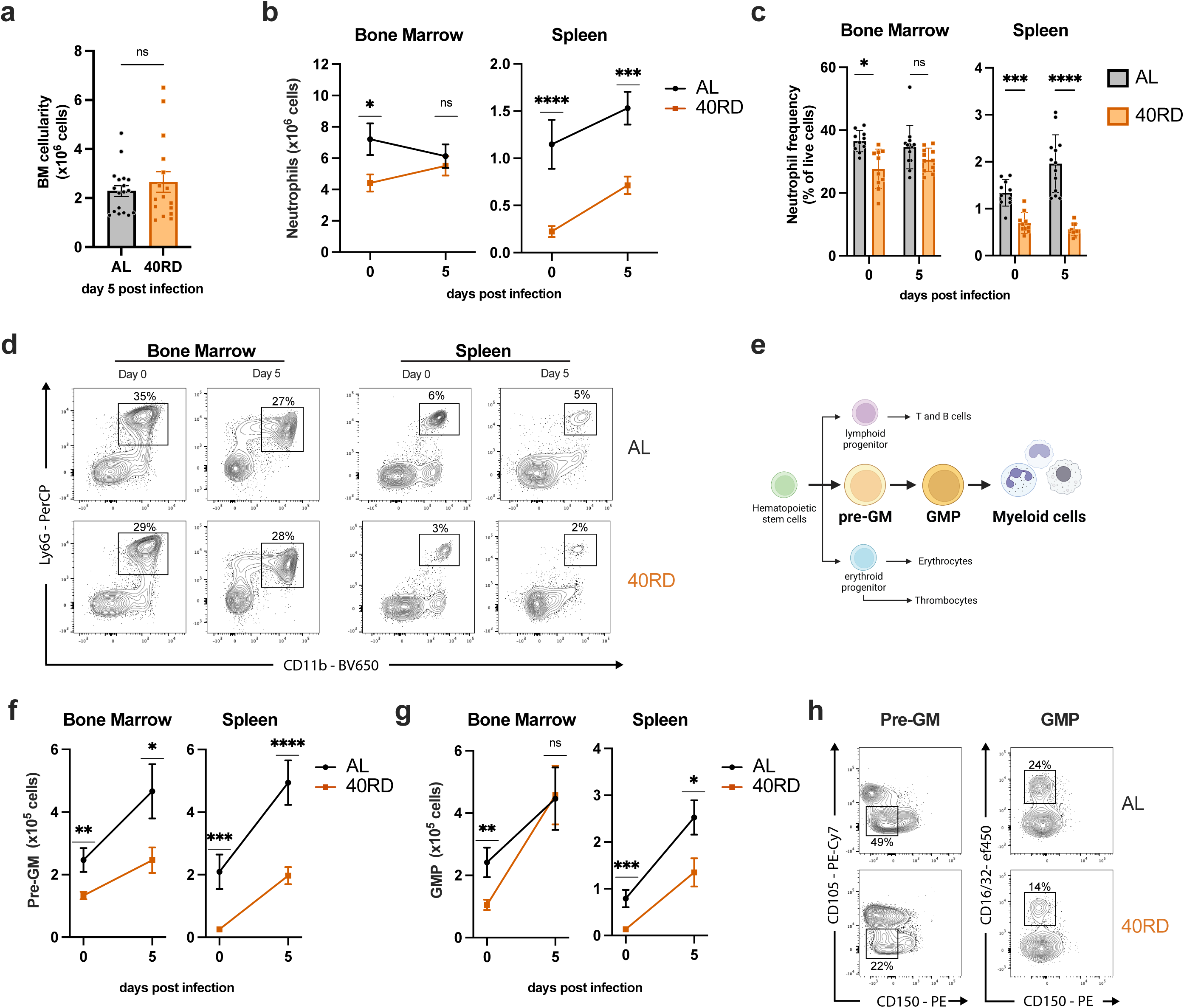
Chronically malnourished mice display dysregulated myelopoiesis. (a) Total bone marrow cell counts from day 5 post-infection AL (n=18) and 40RD (n=16) mice. Bone marrow cells and splenocytes were counted and flow cytometry was performed at days 0 (n=10 for both AL and 40RD) and 5 post Lm infection in AL and 40 RD mice to evaluate (b) the total cell number of neutrophils and (c) the relative frequency of neutrophils among live cells. (d) Representative flow cytometry data for the results in (b,c), with average frequencies shown. (e) A simplified schematic representation of myelopoiesis showing pre-GM and GMP as key progenitors in granulocyte/monocyte lineage. Bone marrow cells and splenocytes were counted and flow cytometry was performed at days 0 and 5 post Lm infection in AL and 40 RD mice to evaluate the total cell number of (f) pre-GM cells (Lineage^−^ Sca1^−^ CD117^+^ CD150^−^ CD16/32^−^ CD105^−^) and (g) GMP cells (Lineage^−^ Sca1^−^ CD117^+^ CD150^−^ CD16/32^+^). (h) Representative flow cytometry results for the myeloid progenitor data in (f,g), with average frequencies shown. Statistics: (a-c,f,g) Plotted as mean ± SEM; statistics done via two-tailed Mann-Whitney test for each time point.

Neutrophil and monocyte production is maintained in the bone marrow during steady-state and through emergency myelopoiesis in both bone marrow and spleen upon infection (Janssen et al., 2016; Kim, 2010; Manz and Boettcher, 2014; Yvan-Charvet and Ng, 2019). We thus evaluated how undernutrition affected the key myeloid progenitor populations, pre-granulocyte/monocytes (pre-GM) and granulocyte/monocyte progenitors (GMP) (Fig. 4e). Consistent with a loss of mature neutrophils and monocytes, we found that myelopoiesis was significantly impaired in undernourished mice. Within the bone marrow, pre-GM and GMP cells were present at lower numbers in steady-state 40RD mice (Fig. 4f,g). Upon infection, these bone marrow progenitor populations were capable of undergoing expansion, with pre-GM cells failing to reach AL levels whereas GMP cells reached comparable numbers (Fig. 4f-h). These defects were more exaggerated in the spleen, where 40RD mice displayed diminished numbers and frequency of pre-GM and GMP cells both pre-and post-infection (Fig. 4f-h). Altogether, we find that undernutrition impairs steady-state myelopoiesis and extramedullary emergency myelopoiesis, blunting the innate immune response against a bacterial pathogen.

### Refeeding intervention reverses wasting, stunting, and global immune atrophy

One of the primary strategies taken to support patients with undernutrition with health complications, including infections, is to refeed them by slowly increasing caloric intake to levels expected for their age group (Ashworth et al., 2003). Despite the widespread employment, the effects of this intervention on immune dysfunction are not known. To address this, we developed a refeeding protocol that safely reintroduces *ad libitum* feeding to 40RD mice to test whether defects in immune responses can be rescued through weight gain and nutrient availability. Undernourished mice undergoing a refeeding protocol (RF) were maintained on a standard 40RD diet for 4 weeks or until 10% BWL. Then, the RF mice were given 10% extra food by weight every two days for a week. On the last day of the refeeding protocol, the RF mice were given *ad libitum* access to feed. Mice were maintained for an additional 6-8 weeks on the *ad libitum* feed until they reached a normal weight range for their age (Fig. 5a). The RF group was able to regain IBW during the refeed period and continued to gain weight at an accelerated pace for the first month (Fig. 5b). After the initial increase in weight, the weight gain pace of RF mice slowed down to the same rate as the AL group (Fig. 5c). We further observed that while RF mice were able to recover growth, the mice maintained a modest but significantly shorter body length than AL controls for their age (Fig. 5d), in keeping with persistent stunting common in patients with undernutrition (Ashworth et al., 2003).

**Figure 5.**
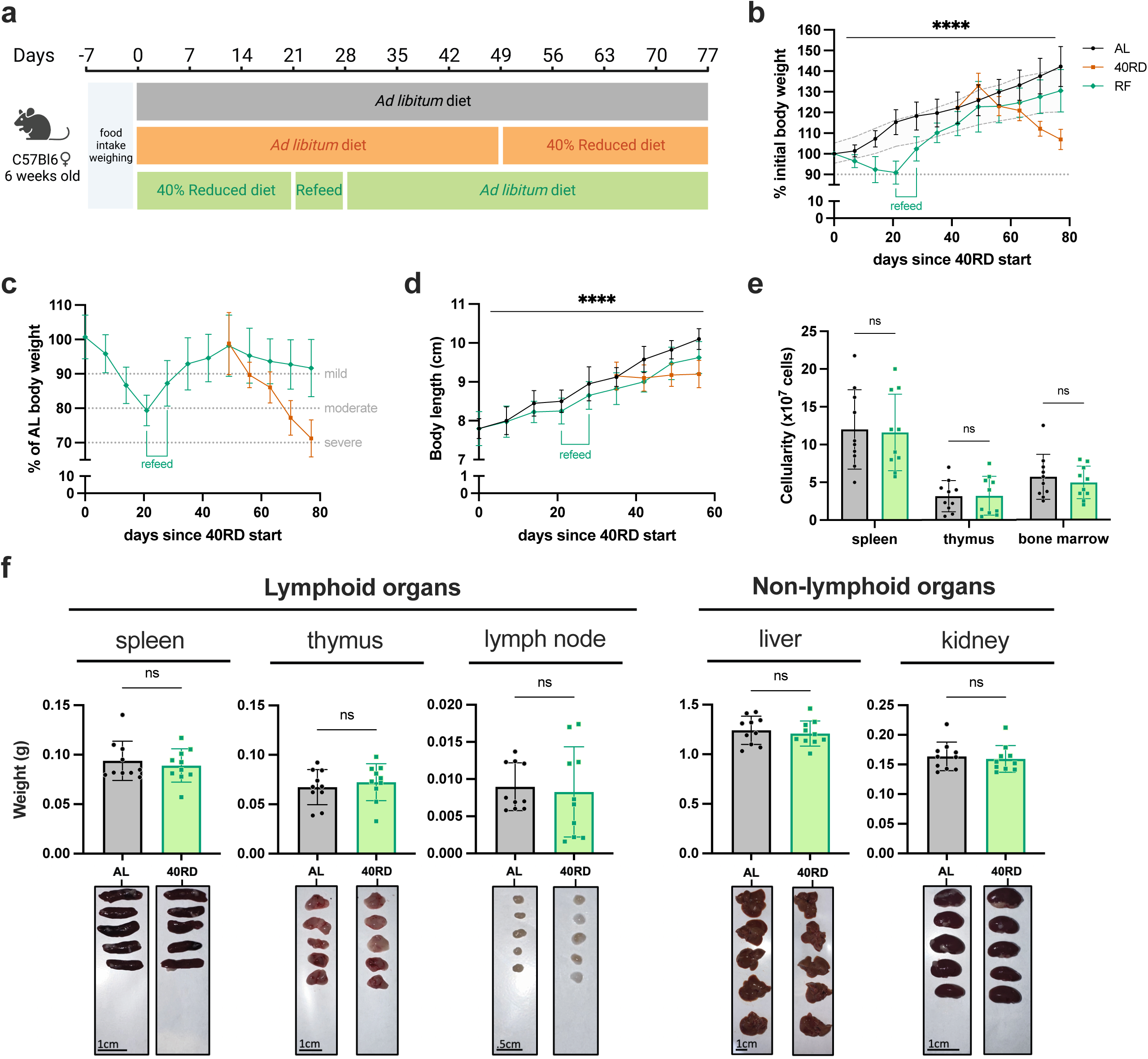
Refeeding intervention reverses wasting, stunting, and global immune atrophy. (a) Schematic of the experimental design for refeeding intervention (RF) in comparison to age-matched 40RD and control AL diet. (b) Body weight of AL (n=25), 40RD (n=10), and RF (n=25) mice as a percentage of their initial body weight over time. The dotted line represents 10% of initial body weight lost, and the shaded area represents the normal weight range for age-matched female C57Bl6 mice(C). (c) Body weight of 40RD (n=10) and RF (n=25) mice as a percentage of age matched average AL body weight over time. Each dotted line represents clinical designations of undernutrition severity. (d) Body length of AL and 40RD mice over time, measured from the nose tip to the base of the tail (n=10). (e) Total cell counts for whole spleen, thymus, and bone marrow from AL and RF mice (n=10). (f) Comparative weights of AL and RF lymphoid and non-lymphoid tissues (n=10) with representative photos of the corresponding organs. Scale bars 1 cm (0.5 cm for lymph nodes). Statistics: (b-f) Plotted as mean ± SD; (b,d) simple linear regression with slope comparisons; and (e,f) two-tailed Mann-Whitney test.

With the refeeding protocol established, we set out to test whether restoring body mass was sufficient to reverse the lymphoid atrophy found in 40RD mice. Paralleling recovery in weight, we found that cellularity in the spleen, thymus, and bone marrow were comparable between AL and RF groups (Fig. 5e). Moreover, we observed that the weight of lymphoid organs has returned to the AL levels in the RF group (Fig. 5f). Together, these findings indicate that refeeding intervention is sufficient to restore global lymphoid atrophy.

### Refeeding intervention fails to restore immunocompetency and normal myelopoiesis

After observing successful reversal of the lymphoid atrophy in RF mice, we tested whether refeeding would be sufficient to restore their ability to control *L. monocytogenes* infection. To test this, we infected age-matched AL, 40RD, and RF mice with a sub-lethal dose of Lm-OVA (Fig. 6a). While refeeding was able to limit morbidity compared to 40RD mice, a portion of RF animals succumbed to infection whereas all AL mice remained viable (Fig. 5b). Consistent with this, we observed that mice with ongoing malnutrition maintained the highest pathogen burdens and that RF mice were unable to resolve infection with the same kinetics as AL mice, with less than half RF mice clearing bacteria (Fig. 5c). These data indicate that despite refeeding being sufficient to reverse malnutrition-induced lymphoid atrophy, prior exposure to chronic undernutrition leads to persistent susceptibility to infection.

**Figure 6.**
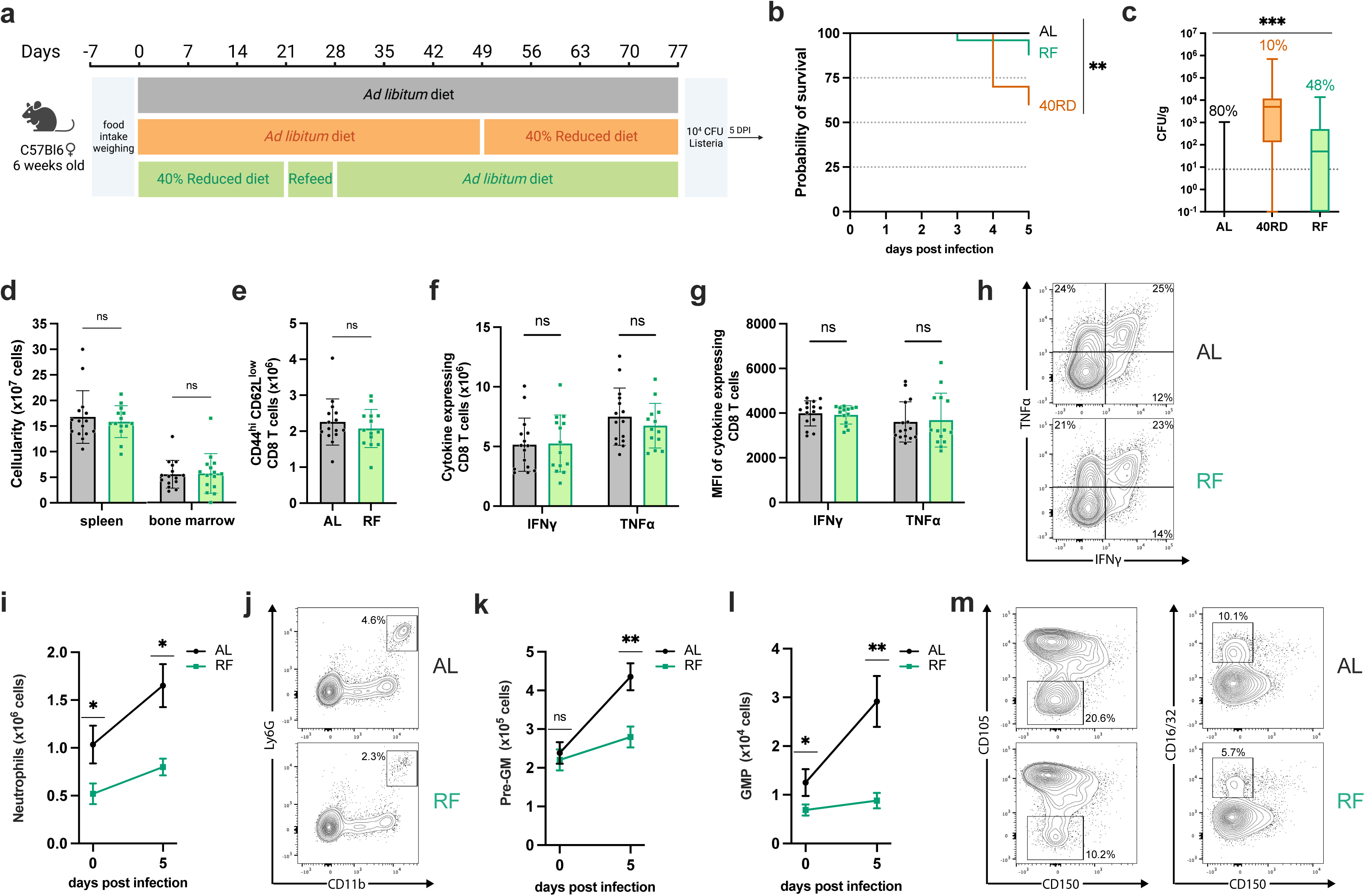
Refeeding intervention fails to restore immunocompetency and normal myelopoiesis. (a) Schematic of Lm infection (10^4^ CFUs per mouse) experimental outline in AL, 40RD, and RF mice. Mice were maintained on the corresponding diet throughout the course of the infection. (b) Probability of survival for infected AL (n=25), 40RD (n=10), and RF (n=25) mice over time. Statistics done via log-rank test. (c) Pathogen burden in liver tissue of 5 DPI AL (n=25), 40RD (n=10), and RF (n=25) mice. Percentage of mice that cleared the pathogen on a given day is represented as numbers above corresponding bars. The dotted line represents the limit of detection. Plotted as box and min to max whiskers; statistics done via Kruskal-Wallis test. (d) Total splenocyte and bone marrow cell counts for AL (n=15) and RF (n=15) mice at day 5 post-infection. (e) Total cell number of CD44^+^CD8^+^ T cells in AL and RF mice at days 0 and 5 post-infection. Splenocytes from AL and RF mice were harvest at day 5 post-infection and stimulated ex vivo with OVA peptide for 6 hours. Intracellular flow cytometry was performed to quantify (f) total cell number of and (g) mean fluorescence intensity (MFI) of antigen-experienced CD8^+^ T cells expressing IFNγ and TNFα. (h) Representative flow cytometry results for data in (f,g), with average frequencies shown. (i) Total splenic neutrophils abundance in AL and RF mice at days 0 and 5 post-infection. (j) Representative flow cytometry results for day 5 post-infection neutrophil data in (i), with average frequencies shown. At days 0 and 5 post-infection, spleens from AL and RF were evaluate for the total number of (k) pre-GM cells and (l) GMP cells. (m) Representative flow cytometry results plots for the myeloid progenitor data in (k,l), with average the average frequencies shown. Statistics: (d-g) Plotted as mean ± SD; statistics done via two-tailed Mann-Whitney test for each category. (i,k,l) Plotted as mean ± SEM; statistics done via two-tailed Mann-Whitney test for each time point.

Having observed this durable impairment in immune protection, we next aimed to identify the specific facets of malnutrition-induced immune dysfunction that failed to recover in RF mice. In keeping with the steady-state recovery observed, post-infection splenic and bone marrow cellularity were equivalent between RF and AL mice (Fig. 6d). Moreover, we found that T and B cell numbers were restored in RF mice and all lymphocyte populations underwent comparable expansion to AL controls after infection (Supplemental 3). Similarly, we observed no differences in the number of antigen-experienced T cells in the spleen nor T cell functional capacity (Fig. 6e-g). Thus, refeeding permits recovery of lymphocyte number, lymphocyte expansion capacity, and T cell function in response to infection, suggesting lymphocyte dysfunction is unlikely to be responsible for the persistent susceptibility observed.

In contrast to the restoration of the lymphocyte response, we observed sustained defects in the myeloid compartment. RF mice displayed impaired peripheral expansion of neutrophil and monocyte populations after infection, both by frequency and number, with neutrophil number diminished in steady-state RF mice as well (Fig 6i,j; Supplemental 4a,b). Within the bone marrow, we observed a reduced frequency in steady-state monocyte and neutrophil abundance as well as reduced total neutrophil numbers, however both populations expanded upon infection, reaching control numbers (Supplemental 4c-f). Having seen this, we next asked whether RF mice exhibited a persistent impairment in either central or emergency myelopoiesis. Whereas myeloid progenitor populations were not perturbed in the bone marrow, we found that RF mice exhibited significantly reduced frequency and numbers of splenic pre-GM and GMP cells following infection, with the GMP population also diminished in steady-state mice (Fig. 6j-l); Supplemental 5). Altogether, our findings demonstrate that refeeding intervention uncouples a global recovery in lymphoid atrophy from an enduring impairment in emergency myelopoiesis, resulting in lasting susceptibility to infection.

## Discussion

Here we employ a murine model of chronic undernutrition that phenocopies many of the hallmarks of human undernutrition (stunting, wasting, and changes in body composition) to characterize how chronic malnutrition leads to nutritionally acquired immunodeficiency. We demonstrate that sustained malnutrition results in global lymphoid atrophy, with loss of both innate and adaptive immune cell populations. Additionally, we identified a unique and severe impairment in neutrophil abundance and myelopoiesis, with a similar but more modest effect on monocytes. Our data suggests that the combination of diminished homeostatic levels of these critical myeloid populations along with impaired emergency myelopoiesis leads to poor control of early *L. monocytogenes* infection, while a blunted T cell response to infection and their premature contraction results in lack of effective infection resolution. We then tested whether a refeeding intervention could restore protective immunity to infection. We found that restoring food access permitted mice to regain weight and size as well as reverse many signs of immunodeficiency. Refeeding was sufficient to recover lymphoid organ atrophy, the abundance of most circulating lymphocytes, and T cell cytokine production. Nonetheless, refed mice continued to display increased susceptibility to *L. monocytogenes* infection along with sustained defects in neutrophil and monocyte abundance and emergency myelopoiesis. These data demonstrate that exposure to chronic undernutrition can result in lasting changes to specific compartments of the immune system, with long-term implications of the health of the individual even after recovering growth and size.

Previous studies have suggested that impaired pathogen responses during undernutrition are driven by lymphocyte loss and functional impairment (Beisel, 1996, 1982; Institute of Medicine, 2000). Low levels of circulating lymphocytes have been well documented in patients with undernutrition and various restrictive diet models (Campbell et al., 2020; Cason et al., 1986; Contreras et al., 2018; Howard et al., 1999; Saha et al., 1977; Schattner et al., 1990; Yang et al., 2009). Our data from the refeeding model suggest that while these defects likely contribute to poor resolution of infection, they are not sufficient to explain why previously malnourished mice remain vulnerable to infection, as these mice recover from lymphoid atrophy and display normal T cell responses. Nonetheless, we find consistent with other groups that undernourished mice exhibit a steady-state reduction in circulating T cells. Upon infection, the T cell compartment is capable of undergoing expansion in malnourished conditions, but the magnitude of this response is blunted and quickly contracts prior to pathogen clearance. We also found that there was reduced abundance of antigen experienced T cells throughout the infection. Additionally, we found that the intrinsic ability of T cells to produce inflammatory cytokines was reduced. In keeping with work demonstrating that T cells from patients with undernutrition display reduced function, we find that T cells from malnourished mice have impaired cytokine producing capacity post-infection. Altogether, our work supports clinical observations and earlier studies delineating the relationship of undernutrition on lymphoid atrophy and T cell dysfunction and has revealed the reversible nature of these defects after refeeding intervention.

In contrast to the recovery found in the adaptive immune cells after refeeding, our work demonstrates that exposure to chronic periods of malnutrition results in durable defects in the myeloid compartment. We find that undernourished mice display impaired myelopoiesis and a loss in abundance in critical peripheral myeloid populations that does not properly recover upon refeeding. Indeed, some small cohort studies have shown that circulating neutrophil numbers are reduced in undernourished children and that hematopoiesis is impacted in anorexic patients (Kohama et al., 2021; Vaisman et al., 2004). Thus, while hematopoiesis and peripheral immune cell numbers are broadly sensitive to an animal’s ongoing nutritional status, specific subsets are uniquely sensitive to periods of undernutrition and exhibit nutritional scarring that uncouples their activity from future improvements in dietary intake. These findings raise the possibility that lasting epigenetic changes within the myeloid compartment occur during exposure to undernutrition, akin to what has been observed for trained immunity (Netea et al., 2020). Indeed, a recent study in a small cohort of undernourished children reported changes in H3 acetylation in peripheral blood mononuclear cells (Kupkova et al., 2021).

While this work provides a novel link between myelopoiesis and nutritional state, many questions remain. Our study investigated the effect of chronic undernutrition during *L. monocytogenes* infection, however the extent to which these findings apply more broadly to other models of bacterial infection or innate inflammation are unclear. Moreover, our malnutrition model broadly restricted food intake without regard to the components of diet that are required for immunocompetency nor consideration for potential interactions of the magnitude of food restriction with immunodeficiency and the rate of onset. It will be of interest in future studies to resolve the extent to which the phenotypes observed are due to the gross loss of calories, specific macronutrients, certain micronutrients, and the grade of dietary restriction enforced. In addition, our study focused on the effects of dietary restriction on adult, normal weight mice. Whether similar phenotypes would emerge if obese animals were dietarily restrict and if weight loss was achieved through pharmacologic means (e.g. GLP-1 targeted drugs) is not clear. Likewise, how similar dietary restriction might affect the immune system if it were initiated in juvenile mice is not addressed here. Mechanistically, additional work is needed to address whether the changes in myelopoiesis are due to changes within the hematopoietic compartment, non-hematopoietic cells, and/or to systemic levels of key cytokines and hormones – such as G-CSF, interferons, and leptin – both in malnourished and refed mice. Finally, the known impact dietary restriction has on microbiota composition and in turn how this might durably impact the immune system is unaccounted for in this work and warrants investigation (Gordon et al., 2012; Blanton et al., 2016).

Altogether our work offers new insight into the contribution of myelopoiesis in nutritionally acquired immunodeficiency and the ability of refeeding intervention to treat these defects. The burden of morbidity and mortality of infections on undernourished individuals is one of the leading global health crises and evidence of impaired vaccine responses in undernourished people is putting in question the world’s ability to protect the most vulnerable populations with its most reliable strategies for prevention (Bhattacharjee et al., 2021; Bhargava, 2016; Bourke et al., 2019, 2016; Dubos, 1955; Ishikawa et al., 2012; Prendergast, 2015; Rice et al., 2000; Saha et al., 1977). We provide evidence that even prior exposure to food scarcity may be sufficient to permanently alter future immune responses. This suggests that the current dietary status of an individual may be insufficient information to understand the impact nutrition has on their immune system. Future work delineating the specific components of diet required to sustain immune health in the face of food scarcity will be critical for developing nutritional interventions or broader food supplementation programs for at risk populations. We believe that this work indicates periods of poverty and food insecurity may be an important factor in patient medical history as well as for formulating global health policy in these areas.

## Materials and Methods

### Mice and diets

All mice were female on a C57Bl6 background maintained at Children’s Hospital of Philadelphia (CHOP) animal facility. Mice were either purchased from Jackson laboratories at 6 weeks of age and acclimated in the CHOP facility for 1 week or bred in house from breeders also obtained from Jackson laboratories. All animal experiments were approved by IACUC, and mice were cared for in accordance with IAC 21-001325 protocol. All 40RD and RF experiments were performed with mice between 6 and 8 weeks of age. Mice were housed in groups of 4-5 animals for all experiments. All experiments were performed on age-matched littermate controls. All mice were maintained on LabDiet 5001 from weaning/arrival to facility.

For the 40RD diet, mice were maintained on *ad libitum* diet until the start of the experiment. One week before the start of the experiment average food intake was measured per 5 mice per day. Average food consumption for all experimental animals between 6 and 8 weeks of age was 4 g/day/mouse. At the start of the experiment, mice were weighed to make sure that all the animals were in the range of ±1.5 g and randomly assigned to an AL or a 40RD group. AL mice were maintained as before with unlimited access to chow. 40RD mice were placed on the 40% reduced diet by weight, resulting in 2.4 g/day/mouse of feed. Consistent weight loss and no competition for food was observed in 40RD mice. 40RD mice lost 10% of their IBW every 4 weeks. If mice lost over 20% of their IBW they were euthanized. During the 40RD diet, experimental animals did not exhibit behavioral or clinical changes and only moderate changed body composition and growth stunting. When all 40RD mice reached 10%-15% IBW loss, they were considered undernourished and utilized for further experiments.

For the RF diet, mice were set up in the identical way to the 40RD diet until they reached 10%-15% IBW loss. At which point they underwent a refeeding intervention developed using Guidelines for the inpatient treatment of severely malnourished children. The refeeding intervention lasted a week and increased the weight of available food to the restricted mice by 10% every 2 days. On the 40RD diet, mice had access to 2.4 g/day/mouse (60% of the initial average food consumption). On day 1 of refeeding intervention, mice had access to 2.8 g/day/mouse (70%). On day 3, they had access to 3.2 g/day/mouse (80%). On day 5, it was 3.6 g/day/mouse (90%). On day 7, mice were given unrestricted food access. RF mice were then maintained on *ad libitum* feed for a month after they entered normal weight range for their age^88^ and their food intake normalized to the AL age-matched controls. At that point mice were considered refed and were utilized for further experiments.

40RD mice and mice on the 1 week-long refeeding intervention were fed daily between 4pm and 6pm to avoid interference with circadian rhythm. Mice on the 40RD and intervention consumed all food provided to them between feeding windows. All mice were observed daily for signs of sickness. All mice were weighed every 2-3 days for 3 times a week total. Food intake for all mice was recorded either daily, if fed daily, or every 2-3 days for 3 times a week total if fed *ad libitum*. Mice were maintained on assigned diets during infection and sepsis experiments. Mice had unlimited access to drinkable water.

### Body length, clinical score, and body condition score (BCS)

For 30 40RD mice and 30 age-matched AL controls body length and BCS were recorded weekly. For 25 RF mice and 25 age-matched AL controls body length were recorded weekly. Body length was recorded using standard centimeter ruler from base of the tail to the tip of the nose. The length was recorded in 0.25cm increments. BCS was recorded using IACUC Pain and Distress Recognition Policy guidelines in a blinded manner by the same observer for all time points. BCS was rated on the scale of 1 – 5, as follows: 1) emaciated, prominent skeletal structure and distinct vertebrae segmentation; 2) underconditioned, segmentation of vertebrae, dorsal pelvic bones palpable; 3) well-conditioned, vertebrae and dorsal pelvis not prominent; 4) overconditioned, spine is a continuous column and vertebrae palpable only with firm pressure; 5) obese, mouse smooth and bulky with bone structure that disappears under flesh and subcutaneous fat (Ullman-Culleré and Foltz, 1999).

Clinical score is a sum of 7 parameters ranked on the scale of 0 - normal to 4 - severely impaired. The scores were measured by the same observer at all time points in a blinded manner. For the measurement, an animal was transferred to a new empty cage and allowed to acclimate for 30 seconds. Then, appearance of the body, appearance of the eyes, level of consciousness, and activity were observed. Then, the cage was tapped twice for auditory stimulus and the animal was touched gently on the back for touch stimulus or gently tipped over if no locomotion was observed. Then, the animal was restrained with standard hold, allowed to acclimate for 30 seconds, and its breathing rate and quality were observed and recorded.

### Listeria preparation and infection

For all the *Listeria monocytogenes* (Lm) infections, the recombinant Lm strain Lm-OVA, kindly provided by the laboratory of Dr. Hao Shen, was used. Lm were grown to early log phase (OD600 = 0.1) in brain heart infusion media (BHI, BD, cat. 237200) at 37°C, washed in PBS (Corning, cat. 21030CV) and diluted to 10^4^ CFUs per 100uL of PBS using our observation that 1 OD unit = 3×10^9^ CFUs. The CFU count was confirmed after infection by plating an aliquot of diluted Lm on BHI plate and incubating overnight at 37°C. Experimental animals were temporarily sedated using Isoflurane (Med Vet International, cat. RXISO-250). 100uL of Lm suspension was injected retro-orbitally per mouse. Mice were observed for full anesthesia recovery. All mice were observed for signs of sickness during infection progression. Any mice passed before the endpoint were tested for CFUs in their liver and recorded as “succumbed to infection” if CFUs were detected or “censored” if CFUs were not detected. For 10 40RD mice and 10 age-matched AL controls, clinical scores were recorded daily at the time of infection and for 14 days following infection.

### Tissue collection and processing

At the end of the experiment, mice were euthanized by carbon dioxide and subsequent cervical dislocation. Either all or a subset of the following tissues were collected, depending on the type of experiment: spleen, bone marrow, thymus, liver, kidney, and inguinal lymph nodes. For steady state assessment, all the tissues were weighed and photographed except for bone marrow before being processed. Liver and kidney were then disposed of and only spleen, thymus, and bone marrow were processed. For the infection model, only spleen and bone marrow were processed, and liver was used for CFU counting described in the next section. Bone marrow was harvested and homogenized by flushing bones with 25G needle and 10mL of cold PBS. Thymus and spleen were harvested in full, homogenized in PBS and strained through a 70-micron cell filter (Celltreat, cat. 229483). All homogenates were resuspended in 1mL of ACK buffer (Quality Biological, cat. 118-156-101) for 2 minutes to perform red blood cell lysis. The reaction was quenched with 10mL of RPMI 1640 supplemented with 10% FBS (Fisher Scientific, cat. BP9703100), 1% Penicillin-streptomycin (Mediatech, cat. MT30-002-CI), 1% L-Glutamate (Mediatech, cat. 1249), 10mM HEPES (Invitrogen, cat. 15630080), 1mM sodium pyruvate (CCS, cat. 1175), and 0.1% 2-Mercaptoethanol (Life technologies, cat. 21985023). Cells then were counted using a hemocytometer and TrypanBlue (Corning, cat. 25-900-CI).

### Liver processing and CFU count

Livers were harvested from Lm infected mice at days 5, 8, and 14-post-infection as described above. Left lateral lobes were collected and weighed. Then, the lobes were homogenized in 1mL of PBS using the Bead Lysis Kit (Next Advance, cat. PINK5E100) and Next Advance Bullet Blender 5E machine for 10 minutes at power 8. The homogenates were serially diluted in PBS between 1:5 and 1:200,000 using serial dilution. The dilutions were plated on BHI plates and incubated at 37oC overnight. A plate with a countable number of CFUs (20 - 100 colonies) was picked per sample. The CFUs were counted by eye and using weight of the lobe and dilution factor, CFUs/g of liver were calculated.

### Flow cytometry and cytokine staining

For surface staining, 4×10^6^ cells from spleen, thymus, and bone marrow homogenates were washed with PBS and stained with a LIVE/DEAD™ Fixable Aqua Dead Cell Stain (Fisher Scientific, cat. L34966) for 15 minutes at 4°C. Then, cells were washed and stained for 20 minutes at 4°C in 2% Rat Serum (StemCell, cat. 13551) and PBS with the following antibodies based on the type of the panel (details below).

#### Innate immune cells

anti-MHC II (eBioscience M5/114.15.2, cat. 48-5321-80, 1:200), anti-CD11b (eBioscience M1/70, cat. 64-0112-82), anti-CD19 (BioLegend 6D5, cat. 115506), anti-Ly6G (Biolegend 1A8, cat. 127616), anti-Ly6C (eBioscience HK1.4, cat. 17-5932-82), anti-Siglec F (eBioscience 1RNM44N, cat. 127616), anti-TCRβ (eBioscience H57-597, cat. 61-5961-82), anti-CD11c (eBioscience N418, cat. 61-5961-82), anti-F4/80 (eBioscience BM8, cat. 17-4801-82, 1:200), anti-NK1.1 (eBioscience PK136, cat. 47-5941-82)

#### Myelopoiesis

anti-C16/32 (eBioscience 93, cat. 14-0161-81), anti-CD3 (eBioscience 145-2c11, cat. 11-0031-85), anti-CD4 (BioLegend GK1.5, cat. 100406), anti-CD8 (Invitrogen 53-6.7, cat. MA1-10303), anti-CD19 (BioLegend 6D5, cat. 115506), anti-CD11b (BioLegend M1/70, cat. 101206), anti-CD11c (eBioscience N418, cat. 11-0114-85), anti-Ter119 (eBioscience TER-110, cat. 11-5921-85), anti-Ly6G/C (eBioscience RB6-8C5, cat. 11-5931-85), anti-B220 (eBioscience RA3-6B2, cat. 11-0452-85), anti-NK1.1 (eBioscience PK136, cat. 11-5941-85), anti-Sca1 (eBioscience D7, cat. 45-5981-82), CD150 (eBioscience mShed 150, cat. 12-1502-82), CD117 (eBioscience 2B8, cat. 61-1171-82), CD105 (eBioscience MJ7/18, cat. 25-1051-82), CD135 (eBioscience A2F10, cat. 17-1351-82), CD48 (BD HM48.1, cat. 561242, 1:200)

#### Effector T cells

anti-CD4 (eBioscience GK1.5, cat. 64-0041-82), anti-CD8 (Invitrogen 53-6.7, cat. MA1-10303), anti-CD44 (eBioscience IM7, cat. 45-0441-82), anti-CD127 (eBioscience A7R34, cat. 12-1271-82, 1:200), anti-KLRG1 (eBioscience 2F1, cat. 25-5893-82)

After staining, cells were washed with PBS. If cells were extracted from infected mice, they were also fixed for 30 minutes at 4°C using eBiosciences Intracellular Fixation Buffer (Termo, cat. 88-8824-00).

For intracellular cytokine staining, 4×10^6^ splenocytes from infected mice were plated in 100uL of supplemented RPMI 1640 described above in U-bottom tissue culture treated 96-well plates (Celltreat, cat. 229190). Cells were stimulated with 2 μg/mL of SIINFEKL peptide (Anaspec, cat. AS-60193-5) and 1 mg/mL of Brefeldin A (Invitrogen, cat. inh-bfa) added 1 hour after the peptide for a total of 6 hours at 37°C. After incubation, cells were washed with PBS and stained with L/D aqua dye for 15 minutes at 4°C. Then, cells were washed and stained for 20 minutes at 4°C in 2% Rat Serum and PBS with the following antibodies: anti-CD4 (eBioscience GK1.5, cat. 64-0041-82), anti-CD8 (Invitrogen 53-6.7, cat. MA1-10303), anti-CD44 (eBioscience IM7, cat. 45-0441-82). After staining, cells were washed with PBS and fixed for 30 minutes at 4°C using eBiosciences Intracellular Fixation Buffer. Cells then were washed using eBiosciences Intracellular Permeabilization Buffer (Termo, cat. 88-8824-00) and stained for 30 minutes at room temperature n 2% Rat Serum and Permeabilization Buffer with the following antibodies: anti-TNFα (eBioscience MP6-XT22, cat. 12-7321-82), anti-IL-2 (eBioscience MQ1-17H12, cat. 25-7029-42, 1:200), anti-IFNγ (eBioscience XMG1.2, cat. 17-7311-82).

All antibodies used at 1:300 dilution unless otherwise specified. Regardless of staining, all cells were washed with PBS at the end of the protocol and resuspended in PBS to perform flow cytometry. Stained samples were analyzed on a 4 laser CytoFlex by Beckman Coulter using an automated plate reader. Data were analyzed using FlowJo 10.5.3 software.

### Statistics and reproducibility

All experiments were repeated at least 3 independent times. Data was represented either as a mean ± standard deviation (SD) or standard error of mean (SEM). For measurements over time within the same experimental cohort, statistical analysis was done via simple linear regression comparing slopes or elevation. For survival curves, log-rank test was performed. For all other pairwise comparisons, Mann-Whitney test was performed. For all other measurements, statistical analysis was done via non-parametric one-way (Kruskal-Wallis) or two-way (mixed-effect) ANOVA depending on the number of variables. P-values are recorded on the graphs as follows: **** for p < 0.0001; *** for p < 0.001; ** for p < 0.01; * for p ≤ 0.05; ns for p > 0.05. All statistical analysis was performed using GraphPad Prism 9.2.0 software.

## Acknowledgments

We thank the Children’s Hospital Flow Cytometry Core for providing support and instrumentation; Drs. Sarah Henrickson, Kelly Jurado, Sunny Shin, and Paula Oliver for feedback and thoughtful discussion; and all members of the Bailis laboratory for providing feedback and support.

## References

Antwi A. Assessment and management of severe malnutrition in children. 2011. West Afr J Med 30:11–8. doi:10.4314/wajm.v30i1.69878.

Ashworth A, Khanum S, Jackson A, Schofield EC. 2003. Guidelines for the inpatient treatment of severely malnourished children. World Health Organization.

Bhattacharjee A, Burr AHP, Overacre-Delgoffe AE, Tometich JT, Yang D, Huckestein BR, Linehan JL, Spencer SP, Hall JA, Harrison OJ, Morais da Fonseca D, Norton EB, Belkaid Y, Hand TW. 2021. Environmental enteric dysfunction induces regulatory T cells that inhibit local CD4+ T cell responses and impair oral vaccine efficacy. Immunity 54:1745–1757.e7. doi:10.1016/j.immuni.2021.07.005

Beisel WR. 1996. Nutrition and Immune Function: Overview. J Nutr 126:2611S–2615S. doi:10.1093/jn/126.suppl_10.2611S

Beisel WR. 1982. Single nutrients and immunity. Am J Clin Nutr 35:417–468. doi:10.1093/ajcn/35.2.417

Bhargava A. 2016a. Undernutrition, nutritionally acquired immunodeficiency, and tuberculosis control. BMJ 355:i5407. doi:10.1136/bmj.i5407

Blanton LV, Barratt MJ, Charbonneau MR, Ahmed T, Gordon JI. 2016. Childhood undernutrition, the gut microbiota, and microbiota-directed therapeutics. Science 352:1533. doi:10.1126/science.aad9359.

Bourke CD, Berkley JA, Prendergast AJ. 2016. Immune Dysfunction as a Cause and Consequence of Malnutrition. Trends Immunol 37:386–398. doi:10.1016/j.it.2016.04.003

Bourke CD, Jones KDJ, Prendergast AJ. 2019. Current understanding of innate immune cell dysfunction in childhood undernutrition. Front Immunol 10. doi:10.3389/fimmu.2019.01728

Campbell C, Marchildon F, Michaels AJ, Takemoto N, van der Veeken J, Schizas M, Pritykin Y, Leslie CS, Intlekofer AM, Cohen P, Rudensky AY. 2020. FXR mediates T cell-intrinsic responses to reduced feeding during infection. Proc Natl Acad Sci USA 117:33446–33454. doi:10.1073/pnas.2020619117

Cason J, Ainley CC, Wolstencroft RA, Norton KR, Thompson RP. 1986. Cell-mediated immunity in anorexia nervosa. Clin Exp Immunol 64:370–375.

Caulfield LE, de Onis M, Blössner M, Black RE. 2004. Undernutrition as an underlying cause of child deaths associated with diarrhea, pneumonia, malaria, and measles. Am J Clin Nutr 80:193–8. doi:10.1093/ajcn/80.1.193.

Cerqueira FM, Kowaltowski AJ. 2010. Commonly adopted caloric restriction protocols often involve malnutrition. Ageing Res Rev 9:424–430. doi:10.1016/J.ARR.2010.05.002

Chatraw JH, Wherry EJ, Ahmed R, Kapasi ZF. 2008. Diminished primary CD8 T cell response to viral infection during protein energy malnutrition in mice is due to changes in microenvironment and low numbers of viral-specific CD8 T cell precursors. J Nutr 138:806–12. doi:10.1093/jn/138.4.806.

Chinen J, Shearer WT. 2010. Secondary immunodeficiencies, including HIV infection. J Allergy Clin Immunol 125:S195–S203. doi:10.1016/j.jaci.2009.08.040

Collins N, Han SJ, Enamorado M, Link VM, Huang B, Moseman EA, Kishton RJ, Shannon JP, Dixit D, Schwab SR, Hickman HD, Restifo NP, McGavern DB, Schwartzberg PL, Belkaid Y. 2019. The Bone Marrow Protects and Optimizes Immunological Memory during Dietary Restriction. Cell. 178:1088–1101.e15. doi: 10.1016/j.cell.2019.07.049

Collins N, Belkaid Y. 2022. Control of immunity via nutritional interventions. Immunity. 55:210–223. doi:10.1016/j.immuni.2022.01.004

Contreras NA, Fontana L, Tosti V, Nikolich-Žugich J. 2018. Calorie restriction induces reversible lymphopenia and lymphoid organ atrophy due to cell redistribution. Geroscience 40:279–291. doi:10.1007/S11357-018-0022-2

Dubos RJ. 1955. Effect of metabolic factors on the susceptibility of albino mice to experimental tuberculosis. J Exp Med 101:59–84. doi:10.1084/jem.101.1.59

Fan Y, Yao Q, Liu Y, Jia T, Zhang J, Jiang E. 2022. Underlying Causes and Co-existence of Malnutrition and Infections: An Exceedingly Common Death Risk in Cancer. Front Nutr 9:814095. doi:10.3389/fnut.2022.814095

Goldberg EL, Romero-Aleshire MJ, Renkema KR, Ventevogel MS, Chew WM, Uhrlaub JL, Smithey MJ, Limesand KH, Sempowski GD, Brooks HL, Nikolich-Žugich J. 2015. Lifespan-extending caloric restriction or mTOR inhibition impair adaptive immunity of old mice by distinct mechanisms. Aging Cell 14:130–138. doi:10.1111/ACEL.12280

Gordon JI, Dewey KG, Mills DA, Medzhitov RM. 2012. The human gut microbiota and undernutrition. Sci Transl Med 4:137ps12. doi:10.1126/scitranslmed.3004347.

Green CL, Lamming DW, Fontana L. 2021. Molecular mechanisms of dietary restriction promoting health and longevity. Nature Reviews Molecular Cell Biology 2021 23:1 23:56–73. doi:10.1038/s41580-021-00411-4

Han SJ, Stacy A, Corral D, Link VM, De Siqueira MK, Chi L, Teijeiro A, Yong DS, Perez-Chaparro PJ, Bouladoux N, Lim AI, Enamorado M, Belkaid Y, Collins N. 2023. Microbiota configuration determines nutritional immune optimization. Proc Natl Acad Sci USA 120:e2304905120. doi:10.1073/pnas.2304905120

Hasegawa A, Iwasaka H, Hagiwara S, Asai N, Nishida T, Noguchi T. 2012. Alternate Day Calorie Restriction Improves Systemic Inflammation in a Mouse Model of Sepsis Induced by Cecal Ligation and Puncture. Journal of Surgical Research 174:136–141. doi:10.1016/J.JSS.2010.11.883

Hess AF. 1932. Diet, Nutrition and Infection. Acta Paediatr 13:206–224. 10.1111/j.1651-2227.1932.tb09220.x

Howard JK, Lord GM, Matarese G, Vendetti S, Ghatei MA, Ritter MA, Lechler RI, Bloom SR. 1999. Leptin protects mice from starvation-induced lymphoid atrophy and increases thymic cellularity in ob/ob mice. J Clin Invest 104:1051–1059. doi:10.1172/JCI6762

Institute of Medicine, 2000. The Role of Nutrition in Maintaining Health in the Nation’s Elderly: Evaluating Coverage of Nutrition Services for the Medicare Population. Washington, DC: The National Academies Press. doi:10.17226/9741

Ishikawa LLW, Da Rosa LC, França TGD, Peres RS, Chiuso-Minicucci F, Zorzella-Pezavento SFG, Sartori A. 2012. Is the BCG vaccine safe for undernourished individuals? Clin Dev Immunol 2012. doi:10.1155/2012/673186

Iyer SS, Chatraw JH, Tan WG, Wherry EJ, Becker TC, Ahmed R, Kapasi ZF. Protein energy malnutrition impairs homeostatic proliferation of memory CD8 T cells. J Immunol 188:77–84. doi:10.4049/jimmunol.

Janssen JW, L BD, V JC, M HP. 2016. Myeloid Cell Turnover and Clearance. Microbiol Spectr 4:4.6.05. doi:10.1128/microbiolspec.MCHD-0005-2015

Jordan S, Tung N, Casanova-Acebes M, Chang C, Cantoni C, Zhang D, Wirtz TH, Naik S, Rose SA, Brocker CN, Gainullina A, Hornburg D, Horng S, Maier BB, Cravedi P, LeRoith D, Gonzalez FJ, Meissner F, Ochando J, Rahman A, Chipuk JE, Artyomov MN, Frenette PS, Piccio L, Berres ML, Gallagher EJ, Merad M. 2019. Dietary intake regulates the circulating inflammatory monocyte pool. Cell 178:1102. doi:10.1016/J.CELL.2019.07.050

Kim CH. 2010. Homeostatic and pathogenic extramedullary hematopoiesis. J Blood Med 1:13–19. doi:10.2147/JBM.S7224

Kohama K, Kotani J, Nakao A. 2021. Effects of Nutrition on Neutrophil Function in Preclinical Studies In: Rajendram R, Preedy VR, Patel VB, editors. Diet and Nutrition in Critical Care. New York, NY: Springer New York. pp. 1–16. doi:10.1007/978-1-4614-8503-2_144-1

Kupkova K, Shetty SJ, Haque R, Petri WA, Auble DT. 2021. Histone H3 lysine 27 acetylation profile undergoes two global shifts in undernourished children and suggests altered one-carbon metabolism. Clin Epigenetics 13:182. doi:10.1186/s13148-021-01173-8

Maleta K. 2006. Undernutrition. Malawi Med J 18:189–205.

Manz MG, Boettcher S. 2014. Emergency granulopoiesis. Nature Reviews Immunology 2014 14:5 14:302–314. doi:10.1038/NRI3660

Martins VJB, Toledo Florêncio TMM, Grillo LP, do Carmo P Franco M, Martins PA, Clemente APG, Santos CDL, de Fatima A Vieira M, Sawaya AL. 2011. Long-lasting effects of undernutrition. Int J Environ Res Public Health 8:1817–1846. doi:10.3390/ijerph8061817

Mbow CC, Rosenzweig LG, Barioni TG, Benton M, Herrero M, Krishnapillai E, Liwenga P, Pradhan MG, Rivera-Ferre T, Sapkota FN, Tubiello YX. 2019. Climate Change and Land: an IPCC special report on climate change, desertification, land degradation, sustainable land management, food security, and greenhouse gas fluxes in terrestrial ecosystems.

Mengheri E, Nobili F, Crocchioni G, Lewis JA. 1992. Protein starvation impairs the ability of activated lymphocytes to produce interferon-gamma. J Interferon Res 12:17–21. doi:10.1089/jir.1992.12.17

Meydani SN, Das SK, Pieper CF, Lewis MR, Klein S, Dixit VD, Gupta AK, Villareal DT, Bhapkar M, Huang M, Fuss PJ, Roberts SB, Holloszy JO, Fontana L. 2016. Long-term moderate calorie restriction inhibits inflammation without impairing cell-mediated immunity: a randomized controlled trial in non-obese humans. Aging 8:1416–1431. doi:10.18632/aging.100994

Murray CJL, Lopez AD, World Health Organization, World Bank, Harvard School of Public Health. 1996. The Global burden of disease : a comprehensive assessment of mortality and disability from diseases, injuries, and risk factors in 1990 and projected to 2020: summary In: Murray CJL, Lopez AD, editors. Global Burden of Disease and Injury Series. Harvard School of Public Health on behalf of the World Health Organization and the World Bank.

Nájera O, González C, Toledo G, López L, Ortiz R. 2004. Flow cytometry study of lymphocyte subsets in malnourished and well-nourished children with bacterial infections. Clin Diagn Lab Immunol 11: 577–80. doi:10.1128/CDLI.11.3.577-580.2004

Netea MG, Domínguez-Andrés J, Barreiro LB, Chavakis T, Divangahi M, Fuchs E, Joosten LAB, van der Meer JWM, Mhlanga MM, Mulder WJM, Riksen NP, Schlitzer A, Schultze JL, Stabell Benn C, Sun JC, Xavier RJ, Latz E. 2020. Defining trained immunity and its role in health and disease. Nat Rev Immunol 20:375–388. doi:10.1038/s41577-020-0285-6

Newsholme A. 1908. The Prevention of Tuberculosis. London: Methuen&Co.

Palma C, La Rocca C, Gigantino V, Aquino G, Piccaro G, Di Silvestre D, Brambilla F, Rossi R, Bonacina F, Lepore MT, Audano M, Mitro N, Botti G, Bruzzaniti S, Fusco C, Procaccini C, De Rosa V, Galgani M, Alviggi C, Puca A, Grassi F, Rezzonico-Jost T, Norata GD, Mauri P, Netea MG, de Candia P, Matarese G. 2021. Caloric Restriction Promotes Immunometabolic Reprogramming Leading to Protection from Tuberculosis. Cell Metab 33:300–318.e12. doi:10.1016/J.CMET.2020.12.016

Pelletier DL. 1994. The Potentiating Effects of Malnutrition on Child Mortality: Epidemiologic Evidence and Policy Implications. Nutr Rev 52:409–415. doi:10.1111/j.1753-4887.1994.tb01376.x

Pena-Cruz V, Reiss CS, McIntosh K. 1989. Sendai virus infection of mice with protein malnutrition. J Virol. 63:3541–4. doi:10.1128/JVI.63.8.3541-3544.1989.

Piccio L, Stark JL, Cross AH. 2008. Chronic calorie restriction attenuates experimental autoimmune encephalomyelitis. J Leukoc Biol 84:940–948. doi:10.1189/JLB.0208133

Prendergast AJ. 2015. Malnutrition and vaccination in developing countries. Philosophical Transactions of the Royal Society B: Biological Sciences 370:20140141. doi:10.1098/rstb.2014.0141

Procaccini C, De Rosa V, Galgani M, Carbone F, Cassano S, Greco D, Qian K, Auvinen P, Calì G, Stallone G, Formisano L, La Cava A, Matarese G. 2012. Leptin-induced mTOR activation defines a specific molecular and transcriptional signature controlling CD4+ effector T cell responses. J Immunol 189:2941–53. doi:10.4049/jimmunol.1200935.

Qiu Z, Khairallah C, Sheridan BS. 2018. Listeria Monocytogenes: A Model Pathogen Continues to Refine Our Knowledge of the CD8 T Cell Response. Pathogens 7:55. doi:10.3390/pathogens7020055

Rice AL, Sacco L, Hyder A, Black RE. 2000. Malnutrition as an underlying cause of childhood deaths associated with infectious diseases in developing countries. Bull World Health Organ 78:1207–1221.

Roberts L. 2017. Nigeria’s invisible crisis. Science 356:18–23. doi:10.1126/science.356.6333.18

Rohr JR, Barrett CB, Civitello DJ, Craft ME, Delius B, DeLeo GA, Hudson PJ, Jouanard N, Nguyen KH, Ostfeld RS, Remais J V, Riveau G, Sokolow SH, Tilman D. 2019. Emerging human infectious diseases and the links to global food production. Nat Sustain 2:445–456. doi:10.1038/s41893-019-0293-3

Saha K, Mehta R, Misra RC, Chaudhury DS, Ray SN. 1977. Undernutrition and Immunity: Smallpox Vaccination in Chronically Starved, Undernourished Subjects and Its Immunologic Evaluation. Scand J Immunol 6:581–589. doi:10.1111/j.1365-3083.1977.tb02136.x

Saucillo DC, Gerriets VA, Sheng J, Rathmell JC, Maciver NJ. 2014. Leptin metabolically licenses T cells for activation to link nutrition and immunity. J Immunol 192:136–44. doi:10.4049/jimmunol.1301158.

Schattner A, Tepper R, Steinbock M, Hahn T, Schoenfeld A. 1990. TNF, interferon-gamma and cell-mediated cytotoxicity in anorexia nervosa; effect of refeeding. J Clin Lab Immunol 32:183–184.

Schlaudecker EP, Steinhoff MC, Moore SR. Interactions of diarrhea, pneumonia, and malnutrition in childhood: recent evidence from developing countries. 2011. Curr Opin Infect Dis 24:496–502. doi:10.1097/QCO.0b013e328349287d.

Sinha P, Lönnroth K, Bhargava A, Heysell SK, Sarkar S, Salgame P, Rudgard W, Boccia D, Van Aartsen D, Hochberg NS. 2021. Food for thought: addressing undernutrition to end tuberculosis. Lancet Infect Dis 21:e318–e325. doi:10.1016/S1473-3099(20)30792-1.

Spadaro O, Youm Y, Shchukina I, Ryu S, Sidorov S, Ravussin A, Nguyen K, Aladyeva E, Predeus AN, Smith SR, Ravussin E, Galban C, Artyomov MN, Dixit VD. 2022. Caloric restriction in humans reveals immunometabolic regulators of health span. Science 375:671–677. 10.1126/science.abg7292

Starr ME, Steele AM, Cohen DA, Saito H. 2016. Short-Term Dietary Restriction Rescues Mice From Lethalabdominal Sepsis and Endotoxemia, and Reduces the Inflammatory/Coagulant Potential of Adipose Tissue. Crit Care Med 44:e509. doi:10.1097/CCM.0000000000001475

Sun D, Muthukumar AR, Lawrence RA, Fernandes G. 2001. Effects of calorie restriction on polymicrobial peritonitis induced by cecum ligation and puncture in young C57BL/6 mice. Clin Diagn Lab Immunol 8:1003–1011. doi:10.1128/CDLI.8.5.1003-1011.2001

Taylor AK, Cao W, Vora KP, De La Cruz J, Shieh WJ, Zaki SR, Katz JM, Sambhara S, Gangappa S. 2013. Protein energy malnutrition decreases immunity and increases susceptibility to influenza infection in mice. J Infect Dis 207:501–10. doi:10.1093/infdis/jis527.

Ullman-Culleré MH, Foltz CJ. 1999. Body condition scoring: a rapid and accurate method for assessing health status in mice. Lab Anim Sci 49:319–23.

UNICEF. 1998. The State of the World’s Children 1998. New York.

UNICEF-WHO-World Bank JME Working Group. 2021. Joint Malnutrition Estimates 2021 – Technical notes on country consultations.

United Nations. 2022. UN Report: Global hunger numbers rose to as many as 828 million in 2021. Rome.

United Nations. 2021. Malnutrition [Fact Sheet].

Vaisman N, Hahn T, Karov Y, Sigler E, Barak Y, Barak V. 2004. Changes in cytokine production and impaired hematopoiesis in patients with anorexia nervosa: The effect of refeeding. Cytokine 26:255–261. doi:10.1016/j.cyto.2004.03.006

Yan X, Imano N, Tamaki K, Sano M, Shinmura K. 2021. The effect of caloric restriction on the increase in senescence-associated T cells and metabolic disorders in aged mice. PLoS One 16. doi:10.1371/JOURNAL.PONE.0252547

Yang H, Youm Y-H, Dixit VD. 2009. Inhibition of Thymic Adipogenesis by Caloric Restriction Is Coupled with Reduction in Age-Related Thymic Involution. J Immunol 183:3040. doi:10.4049/JIMMUNOL.0900562

Yvan-Charvet L, Ng LG. 2019. Granulopoiesis and Neutrophil Homeostasis: A Metabolic, Daily Balancing Act. Trends Immunol 40:598–612. doi:10.1016/J.IT.2019.05.004

Zenewicz LA, Shen H. 2007. Innate and adaptive immune responses to Listeria monocytogenes: a short overview. Microbes Infect 9:1208–1215. doi:10.1016/j.micinf.2007.05.008

