## Supplemental Figures for "Malnutrition drives infection susceptibility and dysregulated myelopoiesis that persists after refeeding intervention"

### Supplemental Figure Legends

**Supplemental 1.**

AL and 40RD mice were infected with $10^4$ CFUs of *Listeria monocytogenes* per mouse. At days 0 and 5 post-infection, spleens were harvested, counted, and analyzed by flow cytometry. (a) The total number of splenic B cells. (b) Relative abundance of B cells as a percentage of all live cells. (c) The total number of splenic CD4 T cells. (d) Relative abundance of CD4 T cells as a percentage of all live cells. (e) The total number of splenic CD8 T cells. (e) Relative abundance of CD8 T cells as a percentage of all live cells.

**Supplemental 2.**

AL and 40RD mice were infected with $10^4$ CFUs of *Listeria monocytogenes* per mouse. At days 0 and 5 post-infection, bone marrow and spleens were harvested, counted, and analyzed by flow cytometry. (a) The total number of bone marrow monocytes. (b) Relative abundance of bone marrow as a percentage of all live cells. (c) The total number of splenic monocytes. (d) Relative abundance of splenic monocytes as a percentage of all live cells. Mice were placed on 40RD and peripheral blood was collected at 1 and 2 weeks post-dietary restriction. Blood was then analyzed to quantify the total number of (e) white blood cells (WBC) and (f) blood neutrophils.

**Supplemental 3.**

AL and RF mice were infected with $10^4$ CFUs of *Listeria monocytogenes* per mouse. At days 0 and 5 post-infection, spleens were harvested, counted, and analyzed by flow cytometry. (a) Relative abundance of B cells as a percentage of all live cells. (b) The total number of splenic B cells. (c) Relative abundance of CD4 T cells as a percentage of all live cells. (d) The total number of splenic CD4 T cells. (e) Relative abundance of CD8 T cells as a percentage of all live cells. (f) The total number of splenic CD8 T cells.

**Supplemental 4.**

AL and RF mice were infected with $10^4$ CFUs of *Listeria monocytogenes* per mouse. At days 0 and 5 post-infection, bone marrow and spleens were harvested, counted, and analyzed by flow cytometry. (a) Relative abundance of splenic monocytes as a percentage of all live cells. (b) The total number of splenic monocytes cells. (c) Relative abundance of bone marrow neutrophils as a percentage of all live cells. (d) The total number of bone marrow neutrophils. (e) Relative abundance of bone marrow monocytes as a percentage of all live cells. (f) The total number of bone marrow monocytes.

**Supplemental 5.**

AL and RF mice were infected with $10^4$ CFUs of *Listeria monocytogenes* per mouse. At days 0 and 5 post-infection, bone marrow was harvested, counted, and analyzed by flow cytometry. (a) Relative abundance of pre-GM cells as a percentage of all live cells. (b) The total number of pre-GM cells. (c) Relative abundance of GMP cells as a percentage of all live cells. (d) The total number of GMP cells.

Supplemental 1

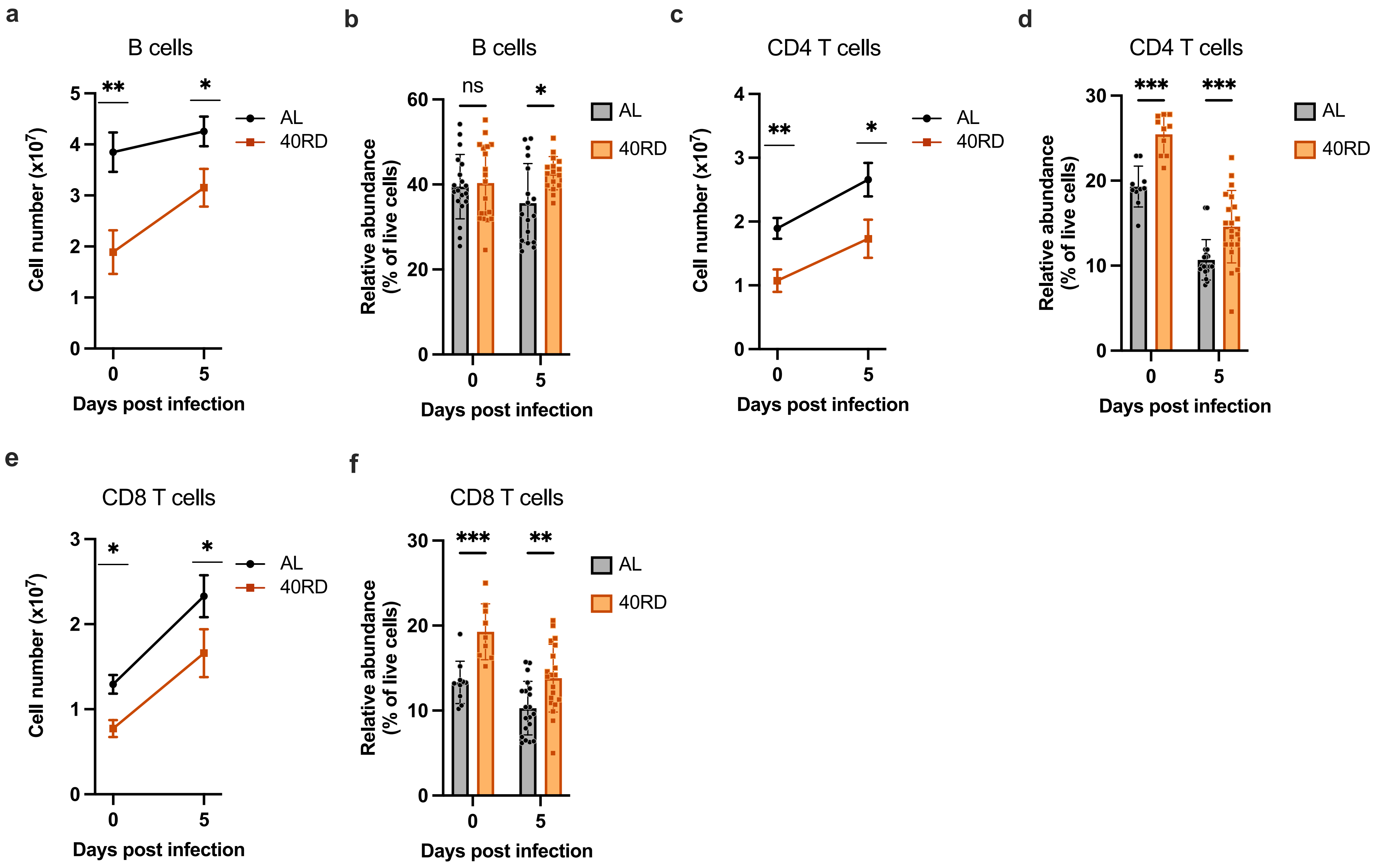

Supplemental 2

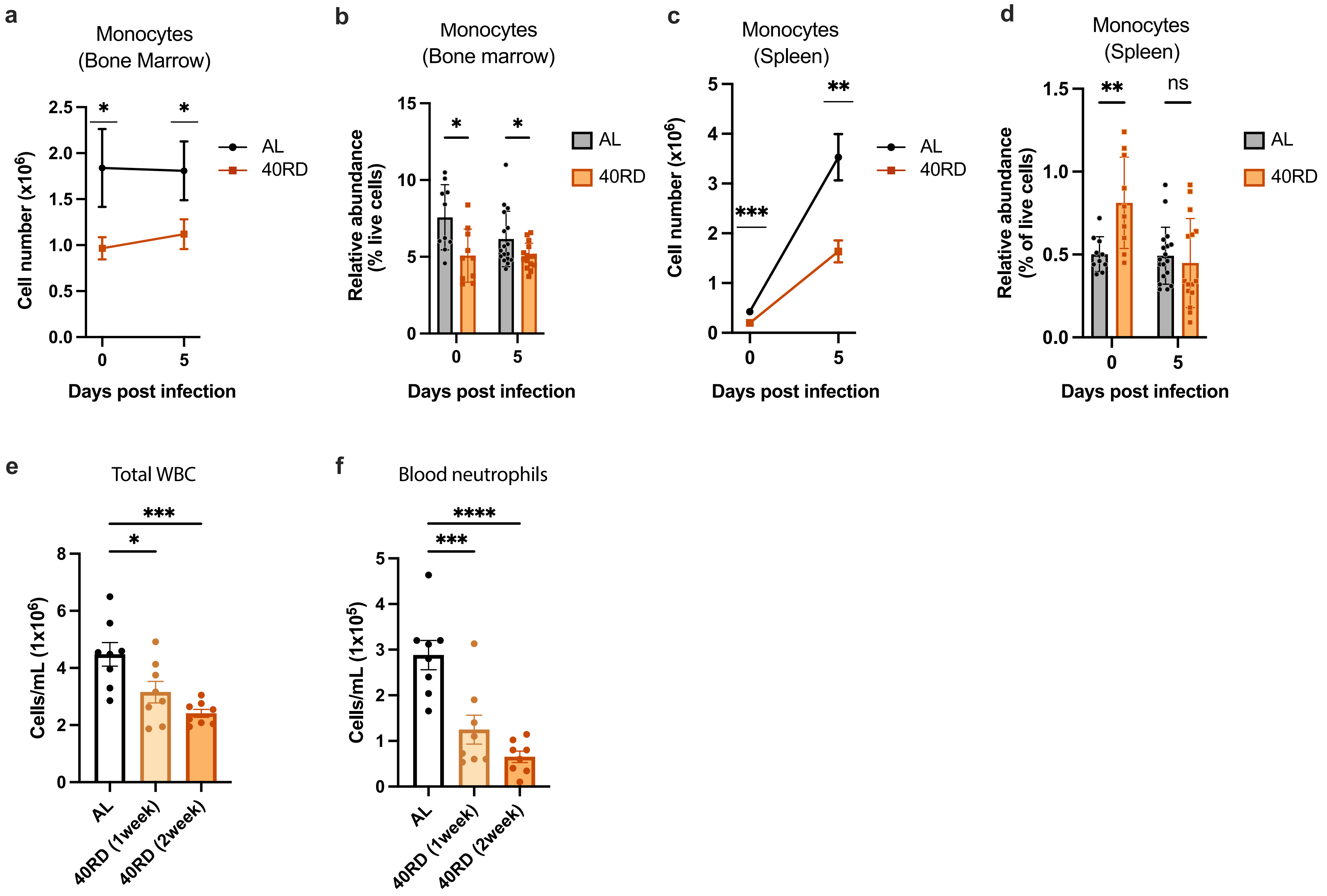

Supplemental 3

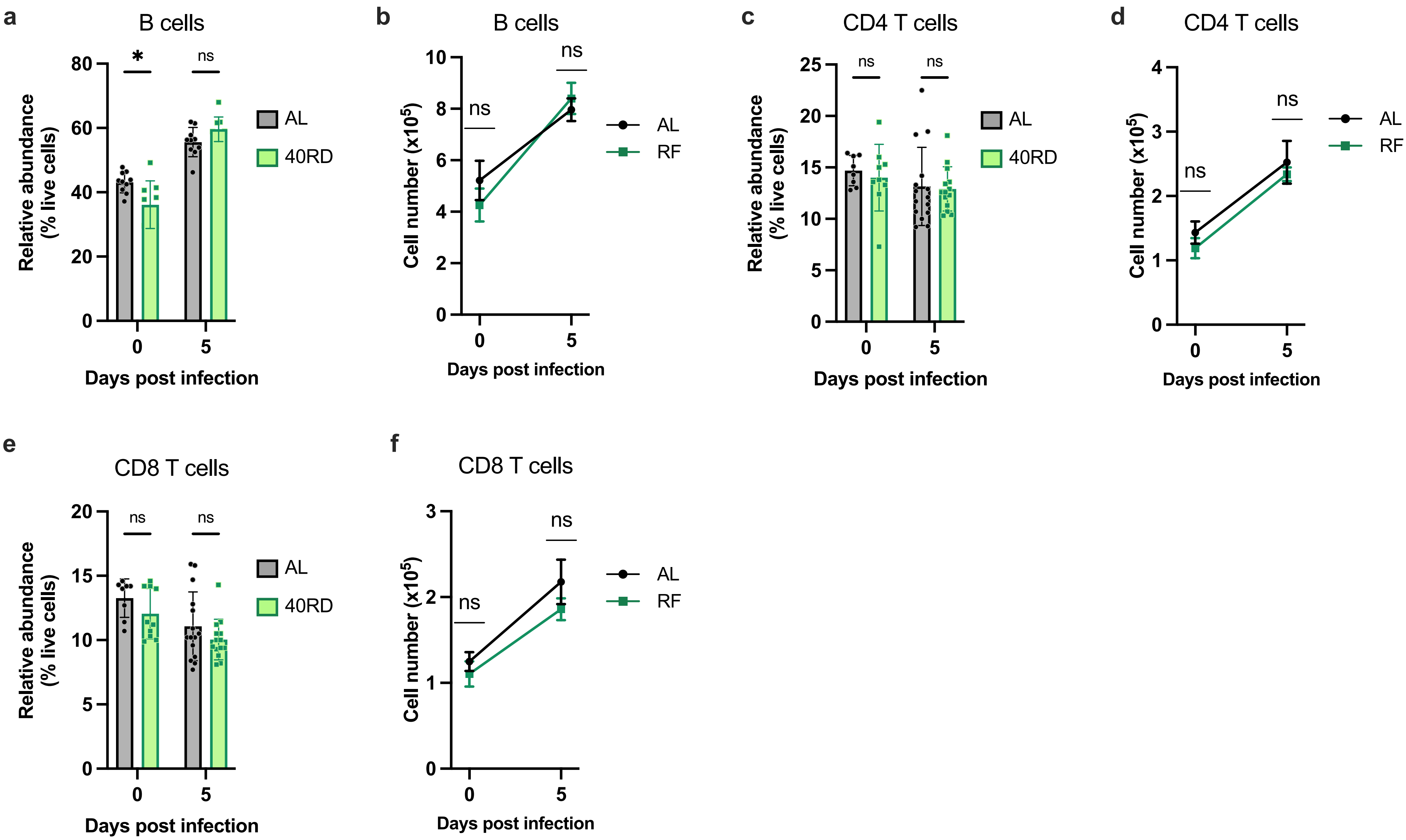

Supplemental 4

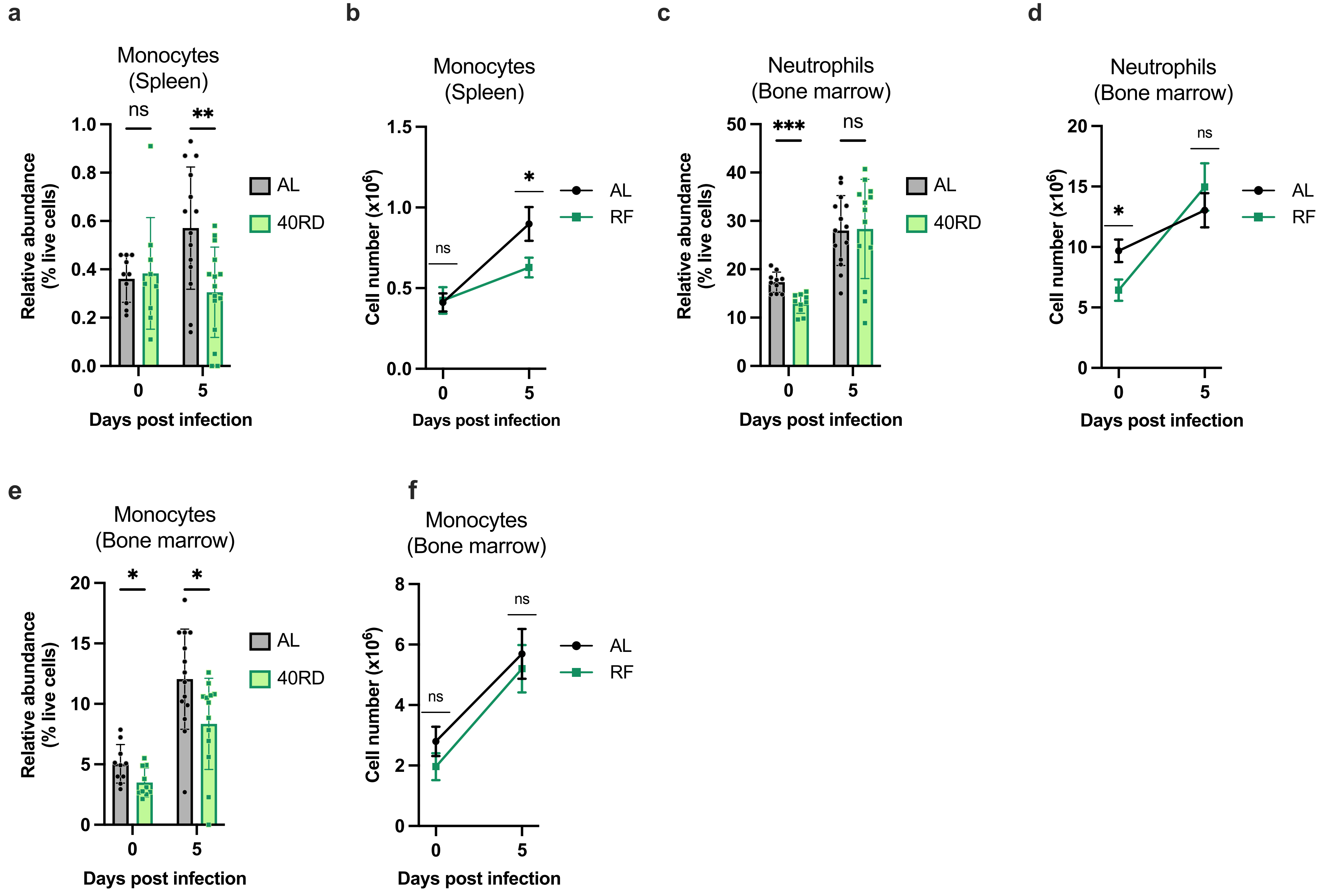

Supplemental 5

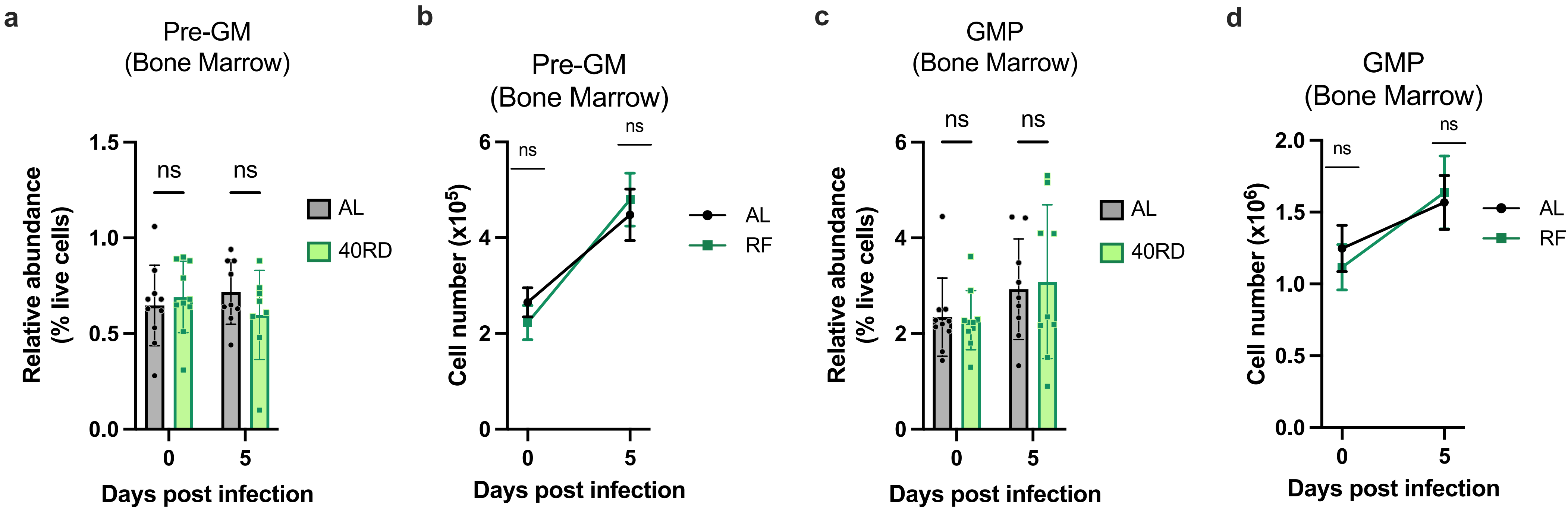
